# Diverse and Location-Specific Roles of PlexinA2, PlexinA4, and NCAM in Developing Hippocampal Mossy Fibers

**DOI:** 10.1101/2024.12.15.628586

**Authors:** Xiao-Feng Zhao, Rafi Kohen, Eljo Y. Van Battum, Ying Zeng, Xiaolu Zhang, Craig N. Johnson, Karen Wang, Brian C. Lim, Juan A. Oses-Prieto, Joshua M. Rasband, Alma L. Burlingame, R. Jeroen Pasterkamp, Matthew N. Rasband, Roman J. Giger

## Abstract

Mossy fibers (MFs) originate from dentate granule cells and innervate area CA3 of the hippocampus. Upon entry of CA3, MFs partition into two prominent axon bundles, the suprapyramidal tract (SPT) and infrapyramidal tract (IPT) and form lamina specific synaptic contacts in the stratum lucidum (SL) and stratum oriens (SO), respectively. Here we employed new mouse lines to dissect the function of Sema6A and its receptors, PlexinA2 (PlxnA2) and PlxnA4, in developing MFs. In *Sema6a^−/−^* mice, MF partitioning into SPT and IPT bundles is incomplete and IPT axons in the SO are overextended, while the SPT correctly innervates the SL. Loss of neuronal *Sema6a* results in defective MF patterning and we show that this involves Sema6A reverse signaling. *Plxna4* controls MF partitioning, SPT axon bundling and laminar targeting to the SL, as well as IPT length. Many of these defects are replicated in mice deficient for PlxnA4 GAP catalytic activity, underscoring the importance of this GAP domain. MFs are tightly fasciculated in *Plxna2^−/−^* mice and fail to separate into SPT and IPT bundles, and defects are significantly reduced in PlxnA2 GAP mutants, highlighting the involvement of GAP-independent signaling events. To further explore the molecular basis of aberrant axon fasciculation, we employed anti-PlxnA2 dependent proximity biotinylation and identified several PlxnA2-associated Ig-CAM family members. We observed a genetic interaction between *Plxna2* and *Ncam1,* but not *Plxna4* and *Ncam1*, for SPT and IPT formation and positioning in CA3. Together, our studies provide insights into the multifaceted and overlapping, yet distinct, functions of PlxnA family members in orchestrating specific guidance decisions in developing MFs.

## Introduction

Proper assembly of complex neural circuits requires the coordinated action of a limited set of axon guidance molecules (*1*). Defects in neuronal connectivity can have far-reaching consequences, contributing to various developmental disorders. Through candidate gene analysis and genome-wide association studies, canonical axon guidance molecules have been identified as candidate risk genes for neuropsychiatric disorders, intellectual disability, and major depressive disorder. For example, mutations in human *PLEXINA2 (PLXNA2)* exhibit variable genetic risks associated with schizophrenia, bipolar disorder, and intellectual disability (*2–4*). Intriguingly, some mutations in *PLXN* family members have been associated with high IQ (*5*). Mutations in genes encoding the PLXNA2 ligands *SEMAPHORIN 5A (SEMA5A*) and *SEMA6A* have been implicated in neuropsychiatric illness, expanding our understanding of the genetic landscape that imparts risk for neuropsychiatric disorder (*6–9*). Collectively, these studies provide a framework for investigating canonical axon guidance molecules in the context of development, cognitive abilities, and brain health.

The Plxns are a family of nine type-1 transmembrane proteins that function as receptors for both membrane-bound and soluble Semas. Within the Plxn family, members can interact directly with Semas, or indirectly using a neuropilin-dependent mechanism (*10*). Plxns can function as canonical Sema receptors (forward signaling) and as ligands for transmembrane Semas, called “reverse signaling” (*11, 12*). Plxns and Semas can interact in *trans* across opposing cell membranes, and in a *cis* configuration within the same cell membrane, thereby influencing the strength of Plxn forward signaling (*12–16*). Members of the immunoglobulin cell adhesion molecule (IgCAM) family function in axon guidance and collaborate with Semas in invertebrates and Sema/neuropilin/Plxn in mammals to ensure proper neural circuits assembly (*17–21*).

Behavioral studies using transgenic mice revealed that loss of *Plxna2* (*22*), or its *Sema5a* (*13*) and *Sema6a* (*6*) ligands, impairs associative learning, sociability, and sensorimotor gating, traits commonly observed in neuropsychiatric disorders. Though studies with global null mutants established important roles for *Plxna2, Plxna4, Sema6a,* and *Sema5a* in cell migration and brain wiring, (*22–29*), our knowledge of downstream signaling mechanisms associated with specific guidance events, and how different guidance receptors collaborate to navigate specific choice points, remains fragmented.

To better understand the contribution of PlxnA2 and PlxnA4 to specific guidance decisions, we focused on hippocampal mossy fiber (MF) wiring. MFs connect the dentate gyrus with area CA3 and form a vital part of the DG-CA3 system, a structure important for memory encoding, storage, and retrieval (*30, 31*). Starting around postnatal day 0 (P0) in rodents, MFs extend radially through the hilus and begin to innervate CA3 (*32*). In the proximal CA3 area, called CA3c, MFs partition to form the SPT and IPT, two prominent fiber bundles travelling immediately above and below the CA3 stratum pyramidales (SP), respectively (**Figure 1A**). SPT axons synapse along the proximal-most region of apical dendrites of pyramidal neurons in CA3 and stop short of the boundary to CA2. IPT axons are comparatively short and preferentially synapse on basal dendrites of CA3 pyramidal neurons. IPT axons extend parallel to the SPT and avoid the SP, except for few axons that cross over to the SP to join the SPT (*33, 34*). In addition, MFs form elaborate connections with GABAergic interneurons in the hilus (Mossy cells), and along the CA3c – CA3a axis (*34–36*). The mature shape of the IPT requires stereotypic pruning of the most distal segment. In the mouse, IPT pruning occurs between P20 and P30 and is regulated, at least in part, by Sema3F/neuropilin-2/Plxn-A signaling (*37–39*).

**Figure 1.**
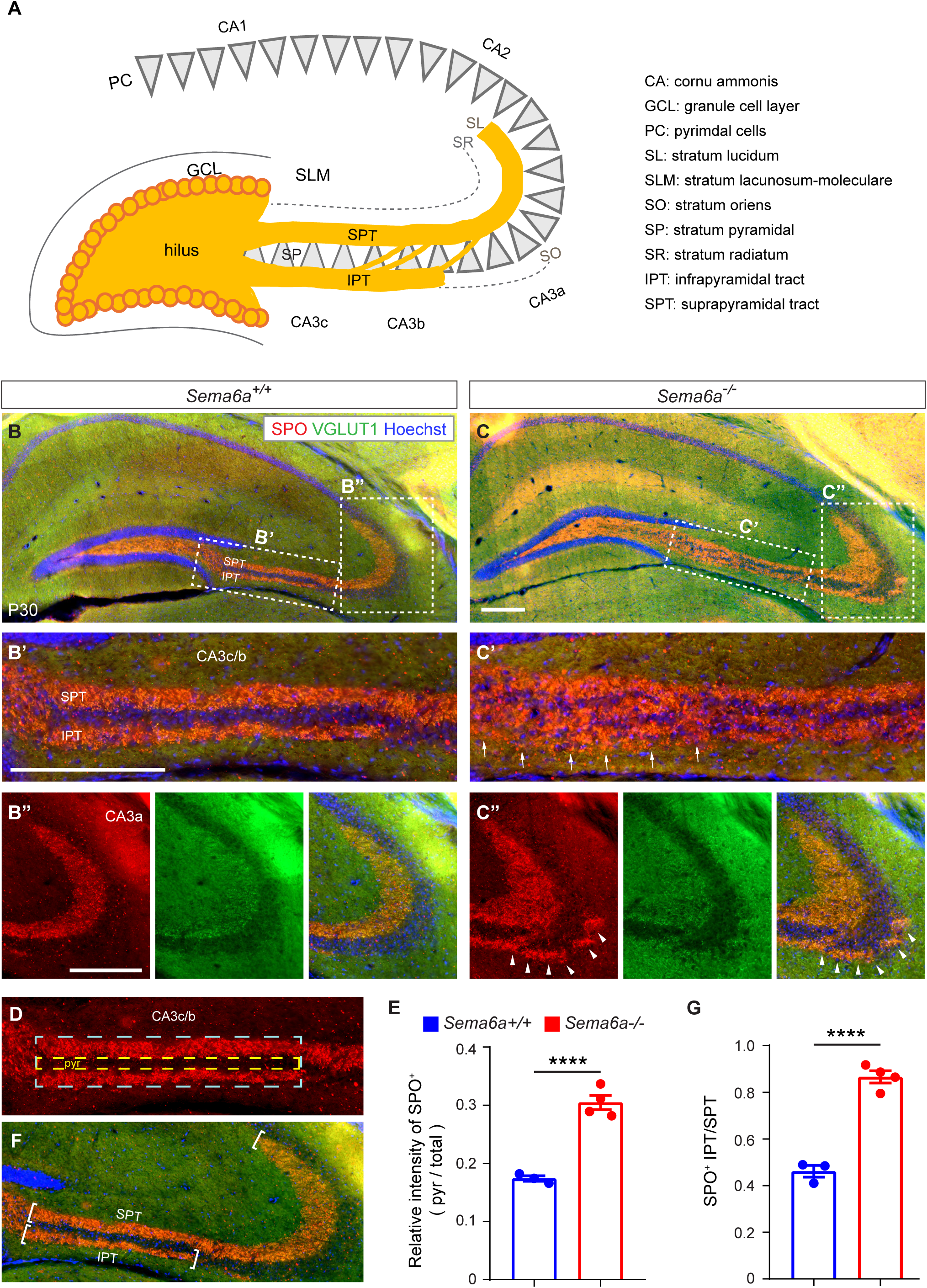
*Sema6a* is necessary for mossy fiber fasciculation and CA3 laminar targeting. (**A**) Schematic of mouse hippocampal MF patterning and laminar targeting along the CA3c-CA3b-CA3a axis. (**B-C’’**) Representative images of coronal brain sections through the dorsal hippocampus of P30 (**B**) *Sema6a^+/+^* control (n=3) and (**C**) *Sema6a^−/−^* (n=4) mice. Sections were stained for the presynaptic makers SPO (red) and VGLUT1 (green) to visualize MF projections. (**B’-C’’**) High magnification images of the CA3c/b and CA3a regions marked with white dotted lines in panels (A) and (B). (B’) Image of CA3c/b region of WT mice. The SPT and IPT are labeled. (C’) Image of CA3c/b region of *Sema6a^−/−^* mice. White arrows indicated region where MF projections fail to partition and bundle into the SPT and IPT. (B’’) Image of WT CA3a region. (C”) Image of CA3a region of *Sema6a^−/−^* mice. Arrowheads indicate overextended IPT axons. Scale bar, 200 µm. (**D, E**) For quantification of defective MF partitioning, the relative intensity of SPO^+^ MF in CA3b/c within the pyramidal cell layer (pyr, area marked with yellow dotted line) over total CA3c/b area (area marked with cyan dotted line) was calculated. (**F**) For quantification of defects in IPT pruning, the length of the IPT and SPT, marked with white brackets, was determined. (**G**) The ratio of the length of the IPT over SPT was calculated. Error bars are SEM. **** p<0.0001, unpaired Student’s *t* test.

Here we pursued a mouse genetic approach to dissect PlxnA2 and PlxnA4 function and signaling mechanisms, aiming at a deeper understanding of how these guidance receptors orchestrate MF patterning *in vivo*. We leveraged conditional gene ablation of *Sema6a* and *Plxna2,* employed a *Sema6a* deletion mutant lacking cytoplasmic sequences (*40*), and developed a PlxnA4 point mutant that harbors a catalytically inactive GAP domain. To probe the extracellular vicinity of PlxnA2 in primary hippocampal neurons, we used antibody directed proximity biotinylation and identified candidate co-receptors, including several IgCAMs. Focusing on neural cell adhesion molecule (NCAM), we show that neuronal *Ncam1* and *Plxna2* genetically interact and are necessary for proper MF separation and bundling into the SPT and IPT in CA3c, as well as IPT pruning, but not for SPT laminar targeting within CA3a. Based on our analysis, we propose a model in which Sema6-PlxnA forward and reverse signaling, in collaboration with NCAM, regulate MF partitioning at the entry of CA3, axon fasciculation into the SPT and IPT, and target zone innervation in CA3b/a. Our studies shed light on the multifaceted and context-dependent functions of guidance molecules in hippocampal circuit assembly.

## Results

### *Sema6a* is necessary for proper development of the infrapyramidal tract

Developing MFs extend radially to establish lamina specific connections with neurons located in the proximal (CA3c), middle (CA3b), and distal (CA3a) areas of CA3, providing opportunities to investigate the underlying axon guidance mechanisms *in vivo* (**Figure 1A**). Previous work has established that Sema6A and Sema6B function as chemorepellents for MFs *in vitro*, governing SPT targeting to the SL *in vivo* (*41*). Analysis of *Sema6a* gene-trap insertion (*Sema6a^lacZ/lacZ^*) mutants did not show any obvious abnormalities in MF laminar targeting (*23*) and *Sema6b^−/−^* mice exhibited mild defasciculation of the SPT and IPT (*41*). However, there is an increase in MF defects in *Sema6a^lacZ/lacZ^*;*Sema6b^−/−^* double mutants, suggestive of potential redundant functions among Sema6 members (*23, 41*).

Because the gene-trap insertion in *Sema6a^lacZ^* occurred at the 3’-end of the sema domain coding sequences (*42*), resulting in a truncated and hypomorphic gene product (*43*), we revisited MF patterning using a recently generated *Sema6a* conditional allele (*Sema6a^flox^* (*43*)). We crossed *Sema6a^flox^* with *CAG-Cre* mice to obtain germline null mutants (*Sema6a^−/−^*). Coronal sections through the dorsal hippocampus of P30 *Sema6a^−/−^* brains, stained with anti-synaptoporin (SPO) and anti-vesicular glutamate transporter 1 (vGLUT1), revealed defects in MF patterning, not previously reported for *Sema6a^lacZ/lacZ^* mice. Specifically, in CA3c/b MFs fail to partition into the SPT and IPT, resulting in aberrant innervation of the SP in CA3c/b (**Figure 1B-C’**). Moreover, in P30 *Sema6a^−/−^* mice IPT axons overshoot their targets in the SO, approaching the apex of the curvature of CA3 (**Figure 1B’’, 1C’’**). To quantify MF fasciculation defects in CA3b/c, we measured SPO immunoreactivity within the SP compared to staining in the SL and SO (*39*). In WT mice, the SPT and IPT are clearly separated and anti-SPO signal within the SP is low (yellow dotted box in **Figure 1D**), compared to total staining intensity (blue dotted box in **Figure 1D**), with a ratio of 0.17 ± 0.01 (n= 3) (**Figure 1E**). This ratio is significantly increased in *Sema6a^−/−^* mice (0.305 ± 0.01, n= 4), indicating defects in SPT and IPT positioning and avoidance of the SP (**Figure 1E**). The lengths of the IPT and SPT were measured (**Figure 1F**) and displayed as the ratio of IPT/SPT, as previously described (*44*). In WT mice, the IPT/SPT ratio is 0.46 ± 0.03 (n= 3) and in *Sema6a^−/−^* mice 0.87 ± 0.03 (n= 4) (**Figure 1G**). Defects in IPT axon length are also observed in 9-month-old mutants (**Figure S1**), indicating that IPT overextension in *Sema6a^−/−^* mice is not resolved at later time points, but persists throughout adulthood. Thus, we report new roles for *Sema6a* in MF development, including partitioning of axons into the SPT and IPT in CA3c and regulation of IPT length in CA3a.

### Complementary, yet overlapping, distribution of PlxnA and Sema6A in the developing hippocampus

In the developing mouse brain, *Plxna2, Plxna4, Sema6a,* and *Sema6b* show highly dynamic patterns of expression (*13, 22, 23, 41*). Longitudinal gene expression analysis in the neocortex, between E10 and P4, revealed expression in excitatory and inhibitory neurons. *Plxna2* expression commences around E12 and *Plxna4* around E14. *Sema6a* is preferentially expressed by *Gad1*^+^ neurons and few excitatory neurons, while *Sema6b* expression begins around E17 and includes excitatory and inhibitory neurons (**Figure S2**).

To assess Sema6A distribution in the early postnatal hippocampus, we took advantage of an epitope tagged *Sema6a* allele (*43*) and explored the relationship to PlxnA2, using double-immunofluorescence labeling. At P1, when granule cells start to extend MFs into the hilus and toward CA3, Sema6A is most prominently found in the stratum lacosum moleculare (SLM) and at reduced levels in the SR and SL (**Figure S3A-D**). At P1, highest levels of PlxnA2 are observed in the hilus and a thin band above the SP along the entire CA3-CA1 region, including the SL, the target zone of SPT axons. In addition, elevated PlxnA2 labeling is observed below the SP in the SO, where IPT axons travel (**Figure S3E-H**). Thus, Sema6A distribution is largely complementary to PlxnA2, with some overlap in the SL. Later, at P7, when MFs have reached their target zones in CA3, PlxnA2 levels are reduced in the SL, but are increased in the SP. At P7, HA-Sema6A is detected in the SL, and levels in the SP are increased compared with P1 (**Figure S3I-L; Figure S4A,B**).

### *Neuronal Sema6a* is necessary for proper MF development

*Sema6a* is strongly expressed by cells in the OL lineage (*45*). Double immunofluorescence labeling of P7 brain sections with anti-HA and anti-MAG or anti-MBP demonstrates presence of HA-Sema6A in myelinating OLs (**Figure S4C-4F**) and little overlap with PDGFRα^+^ OL progenitors or GFAP^+^ radial glia and astrocytes (**Figure S4G-J**). To determine the cell autonomy of MF defects observed in *Sema6a^−/−^* mice, we generated *Sema6a^flox/flox^;Syn1-Cre^+/−^* and *Sema6a^flox/flox^;Olig2-Cre^+/−^* conditional mutants (**Figure 2**). In P30 *Sema6a^flox/flox^;Olig2-cre^+/−^* brains, MF develop normally and are indistinguishable from parallel processed *Sema6a^flox/flox^* controls (**Figure 2A-2B’’**). However, in P30 *Sema6a^flox/flox^;Syn1-cre^+/−^* brains, MF partitioning into the SPT and IPT is impaired, and axons show aberrant innervation of the SP (**Figure 2C, 2C’, and 2D**). Moreover, in *Sema6a^flox/flox^;Syn1-cre^+/−^* mice, the IPT is overextended. While defects in *Sema6a^flox/flox^;Syn1-cre^+/−^* mice resemble those in *Sema6a^−/−^* mutants, they are less pronounced, possibility due to incomplete cre recombination or the contribution of Sema6A produced by non-neuronal cell types (**Figure 2C’’ and 2E**). Together, this suggests that *Sema6a* functions in neurons, not OLs, and that it is important for proper MF patterning *in vivo*.

**Figure 2.**
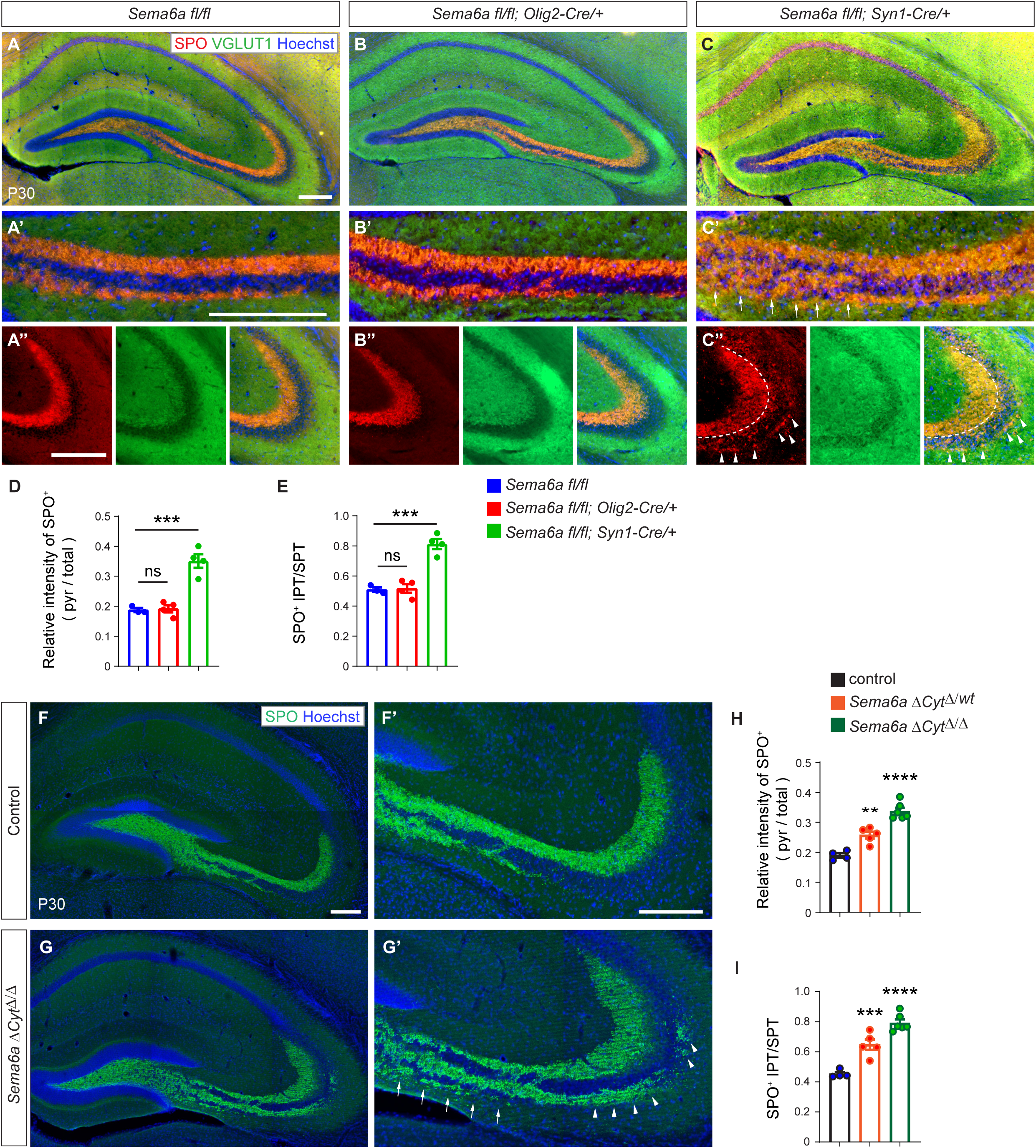
Neuronal Sema6A and Sema6A reverse signaling participate in MF patterning. (**A-C’’**) Representative images of coronal sections through the dorsal hippocampus of P30 brains stained for the presynaptic markers SPO and vGLUT1 to visualize MF projections in (**A**) *Sema6a^fl/fl^* controls (n=3), (**B**) *Sema6a ^fl/fl^;Olig2-Cre/+* (n=4), and (**C**) *Sema6a ^fl/fl^;Syn1-Cre/+* (n=4) mice. (**A’** - **C’**) Higher magnification images of area CA3c/b shown in A-C. (C’) White arrows demarcate the region of defective MF partitioning in *Sema6a ^fl/fl^; Syn1-Cre/+* mice. (**A’’** - **C’’**) Higher magnification images of area CA3a. (C’’) Arrowheads in CA3a highlight extension of IPT axons in *Sema6a ^fl/fl^; Syn1-Cre/+* mice. Scale bar, 200 µm. (**D**) Quantification of defective MF partitioning in area CA3b/c. Shown is the ratio of SPO^+^ MF in the pyramidal cell layer/total. (**E**) Quantification of aberrant IPT length. The ratio of IPT/SPT length is shown. Error bars are SEM. *** p<0.001. ns, not significantly, one-way ANOVA. (**F** and **G**) Representative images of coronal sections through the dorsal hippocampus of P30 brains of control (*Sema6a+/+*) and *Sema6aΔcyt^Δ^*^/*Δ*^ mice, stained with anti-SPO and Hoechst nuclear dye (**F’** and **G’**) Higher magnification of CA3 regions shown in F and G. Defects in MF partitioning are indicated with arrows and defects in IPT length with arrow heads. Scale bar, 200 µm (**H**). Quantification of defective MF partitioning in area CA3c/b in control, n= 4, *Sema6aΔcyt^Δ/wt^* n= 5, and *Sema6aΔcyt^Δ^*^/*Δ*^ n= 6 mice. Shown is the ratio of SPO^+^ MF in the pyramidal cell layer/total. (**I**) Quantification of aberrant IPT length. The ratio of IPT/SPT length is shown. Error bars are SEM. ** p<0.01, *** p<0.001 and **** p<0.0001, one-way ANOVA.

### *Sema6a* reverse signaling participates in MF development

Sema6A is a canonical ligand for PlxnA2 and PlxnA4 (*46, 47*), but it can also operate as a PlxnA receptor in reverse signaling (*25, 46*). Moreover, Sema6A has been shown to associate with PlxnA in *cis* configuration, thereby attenuating ligand interactions in *trans* (*12, 14*). Because of these different signaling modes, we wished to determine how Sema6A contributes to MF patterning. We took advantage of a recently generated *Sema6a* mouse line (*Sema6aΔcyt*), deficient for exon 19, which encodes the *Sema6a* cytoplasmic region. Because the Sema6A ectodomain remains intact and attached to the plasma membrane in *Sema6aΔcyt* mice, forward signaling and *cis* inhibition are not disrupted. Analysis of MFs at P30, using anti-SPO and anti-vGlut1 immunofluorescence labeling, revealed gene dosage-dependent defects in SPT and IPT partitioning in CA3a. While discrete bundles are formed in *Sema6aΔcyt^Δ/wt^* and *Sema6aΔcyt^Δ^*^/*Δ*^ mice, more axons are found in the IPT compared to the SPT, and the IPT is ectopically positioned in the SP (**Figure 2F-I**). Moreover, in both, *Sema6aΔcyt^Δ/wt^* and *Sema6aΔcyt^Δ^*^/*Δ*^ mice, the IPT is significantly extended when compared to WT mice processed in parallel, indicating that Sema6A reverse signaling is sensitive to gene dosage (**Figure 2F-I**). Together, these studies indicate that some, but not all, MF defects observed in *Sema6a^−/−^* are replicated in *Sema6aΔcyt^Δ/wt^* and *Sema6aΔcyt^Δ^*^/*Δ*^ mutants, highlighting the importance of both, Sema6A forward and reverse signaling, for proper MF development.

### PlxnA4 GAP dependent and independent guidance mechanisms orchestrate DG-CA3 connectivity

To better understand the role of PlxnA4 GAP enzymatic activity in MF patterning, we generated a *Plxna4* variant *Plnxa4(R1745A)*, hereafter referred to as *Plxna4^R^*, in which the catalytically essential arginine 1745 is replaced by alanine. To validate functionally disruption of GAP activity, expression plasmids encoding *Plxna4, Plxna2,* or *Plxna4^R^* were used to transfect HEK293T cells which were then probed for Sema6A ectodomain (Sema6A-Fc) cell surface binding. Cultures were then assayed for alteration in cell shape, including rounding up of cells that support Sema6A-Fc binding. Recombinant PlxnA2, PlxnA4, and PlxnA4^R^ are localized to the cell surface where they strongly support binding of bath applied Sema6A-Fc (**Figure S5A-B**). Importantly, no changes in cell shape were observed upon Sema6A-Fc binding to PlxnA4^R^ (**Figure S5B**). Consistent with previous studies, Sema6A- Fc treatment of cells expressing PlxnA2 or PlxnA4 caused cell rounding (**Figure S5B**). These studies indicate that GAP catalytic activity of PlxnA4 is necessary for Sema6A signaling *in vitro*. Co-expression of Sema6A in *cis* with PlxnA2, PlxnA4, or PlxnA4^R^ greatly reduced Sema6A-Fc binding in *trans* (**Figure S5B**). Together these experiments show that recombinant Plxn4A^R^ is localized to the cell surface, interacts with Sema6A both in *cis* and in *trans* configurations, but is defective for Sema6A triggered signaling pathways that cause changes in cell shape.

To assess the role of PlxnA4 GAP catalytic activity in MF patterning *in vivo,* we employed CRISPR/Cas9 mediated gene editing to introduce the R1745A mutation in the endogenous *Plxna4* gene. Four founder lines were obtained that carry the *Plxna4^R^* allele. Two independent lines were established and bred to homozygosity, resulting in *Plxna4^R/R^* mice, and these were analyzed separately. Introduction of this point mutation did not impact viability or gross morphology (data not shown). Moreover, PlxnA4 protein levels were comparable to WT mice, as assessed by Western blot analysis of P7 brains collected from *Plxna4^+/+^, Plxna4^R/+^,* and *Plxna4^R/R^* pups (**Figure S5C**).

Because PlxnA4 is abundantly expressed in the developing and adult mouse brain, we used anti-PlxnA4 immunofluorescence to stain P30 *Plxna4^+/+^*, *Plxna4^R/R^*, and *Plxna4^−/−^* hippocampal sections (**Figure 3**). The dentate molecular layer, hilus, and MFs are heavily decorated by anti-PlxnA4 in *Plxna4^+/+^* and *Plxna4^R/R^* mice, while *Plxna4^−/−^* brains processed in parallel show no labeling (**Figure 3A-C**). This demonstrates that loss of GAP catalytic activity does not alter PlxnA4 abundance or axonal localization *in vivo*. Importantly, *Plxna4^R/R^* mutants show defects in MF patterning, mimicking some, but not all, defects observed in *Plxna4^−/−^* mice. Specifically, in *Plxna4^R/R^* mice, SPT axons are defasciculated in CA3a and no longer confined to the SL, but instead they ectopically innervate the SR, comparable to defects observed in *Plxna4^−/−^* mice (**Figure 3C, 3F-F’**). In both, *Plxna4^R/R^* and *Plxna4^−/−^* mice, IPT axons overshoot their target in the SO, extending toward the apex of the CA3 curvature (**Figure 3C-F’**). Proper partitioning of MFs into SPT and IPT bundles in CA3c is observed only in *Plxna4^R/R^* mice, but not in *Plxna4^−/−^* mice (**Figure 3E’, F’, G, H**). In *Plxna4^−/−^* mice, MFs are severely defasciculated in CA3c and fail to separate in SPT and IPT bundles, resulting in aberrant innervation of the SP (**Figure 3E, E’, G, H**). Thus, PlxnA4 forward signaling requiring GAP catalytic activity is necessary only for SPT laminar targeting to the SL and regulation of IPT length, but not for partitioning of the SPT and IPT in CA3c.

**Figure 3.**
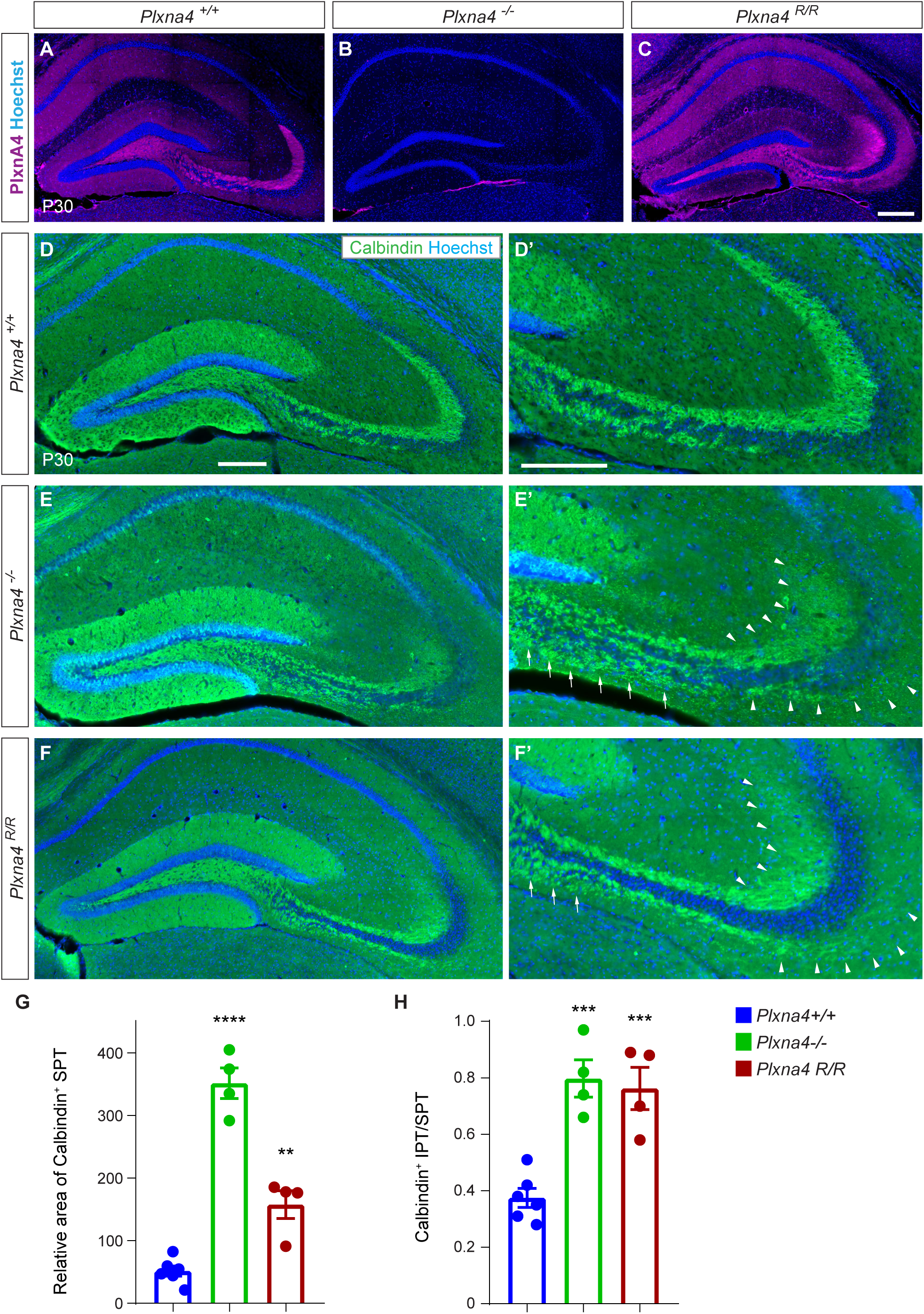
PlxnA4 GAP activity is required for SPT and IPT bundling and target innervation. (**A-C’**) Representative images of coronal sections through the dorsal hippocampus of P30 (A) *Plxna4^+/+^*, (B) *Plxna4^−/−^*, and (C) *Plxna4^R/R^* brains, stained with anti-PlxnA4 and Hoechst nuclear dye. (**D-F**) Images of coronal brain sections through the dorsal hippocampus of P30 (A) *Plxna4^+/+^* (n=6), (B) *Plxna4^−/−^* (n=4) and (C) *Plxna4^R/R^* (n=4) mice, stained with anti-calbindin. (**D’-F’**) Higher magnification images of D-F. Arrows in area CA3c/b and arrowheads in CA3a point to defective MF patterning in *Plxna4^−/−^* and *Plxna4 ^R/R^* mice, respectively. Scale bar for all images, 200 µm. (**G**) Quantification of defective MF partitioning in area CA3c/b. Shown is the ratio of SPO^+^ MF in the pyramidal cell layer/total. (**H**) Quantification of aberrant IPT length. The ratio of IPT/SPT length is shown. Error bars are in SEM. ** p<0.01, *** p<0.001 **** p<0.0001, one-way ANOVA.

Because MF projections to the SPT and IPT fail to separate in *Plxna4^−/−^* mice, it is not clear whether aberrant MF axons in the SO represent SPT axons that have wandered astray and crossed over the SP, or IPT axons that overshoot their target zone in the SO. Because SPT and IPT partitioning still occurs in *Plxna4^R/R^* mice, yet IPT axons overshoot their target, this provides additional evidence that PlxnA4 forward signaling through GAP catalytic activity is necessary for the regulation of IPT axon length.

### PlxnA2 GAP activity is necessary for proper MF patterning

*Plxna2* germline null mice (*Plxna2^−/−^*) show severe defects in MF partitioning into SPT and IPT as well as laminar targeting to the SL and SO. In CA3c/b MFs are tightly fasciculated and pushed ventrally, below the SP. More distally in CA3a, MF projections do not enter the SL and aberrantly innervate the SP (*22, 23*). MF defects are much less pronounced following selective disruption of the PlxnA2 GAP catalytic activity. As previously shown, proper SPT laminar targeting to the SL still occurs in *Plxna2^R/R^* mice (*22*). However, similar to *Plxna2^−/−^* mutants, *Plxna2^R/R^* mice show defects in IPT and SPT partitioning at the entry of CA3 indicative of aberrant axon-axon adhesion (**Figure 5A**). To investigate the contribution of *Sema6a* to defects observed in *Plxna2^−/−^* and *Plxna2^R/R^* mice, we generated compound mutants. In *Sema6a^+/−^*;*Plxna2^+/+^* mice, MF patterning and target innervation along the CA3c-CA3a axis appears largely normal (**Figure 4A-A’’**). In *Sema6a^+/+^;Plxna2^+/−^* mice, separation of the SPT and IPT in CA3c/b is defective, while SPT targeting to the SL in CA3a is largely normal and the IPT is not significantly longer (**Figure 4B-B”**, **Figure 5E).** Analysis of *Sema6a^+/−^*;*Plxna2^+/−^* compound heterozygous mice revealed defects in MF partitioning reminiscent of *Plxna2^+/−^* mice (**Figure 4B-B’** and **Figure 4C-C’’**). In compound null mutants (*Plxna2^−/−^;Sema6a^−/−^*), MF develop largely normal; the SPT and IPT separate at the entry into CA3 and innervation of laminar targets, the SL and SO, is indistinguishable from WT mice (**Figure 4D-D’’**), however the IPT remains overextended (quantification shown in **Figures 5D, E**). This shows that aberrant MF bundling, and innervation of the SP observed in *Plxna2^−/−^* mice requires *Sema6a* function. Conversely, defects in SPT and IPT separation in *Sema6a−/−* mice require *Plxna2* function. However, the overextension of IPT axons observed in *Sema6a−/−* mice occur independently of whether *Plxna2* is present or absent.

**Figure 4.**
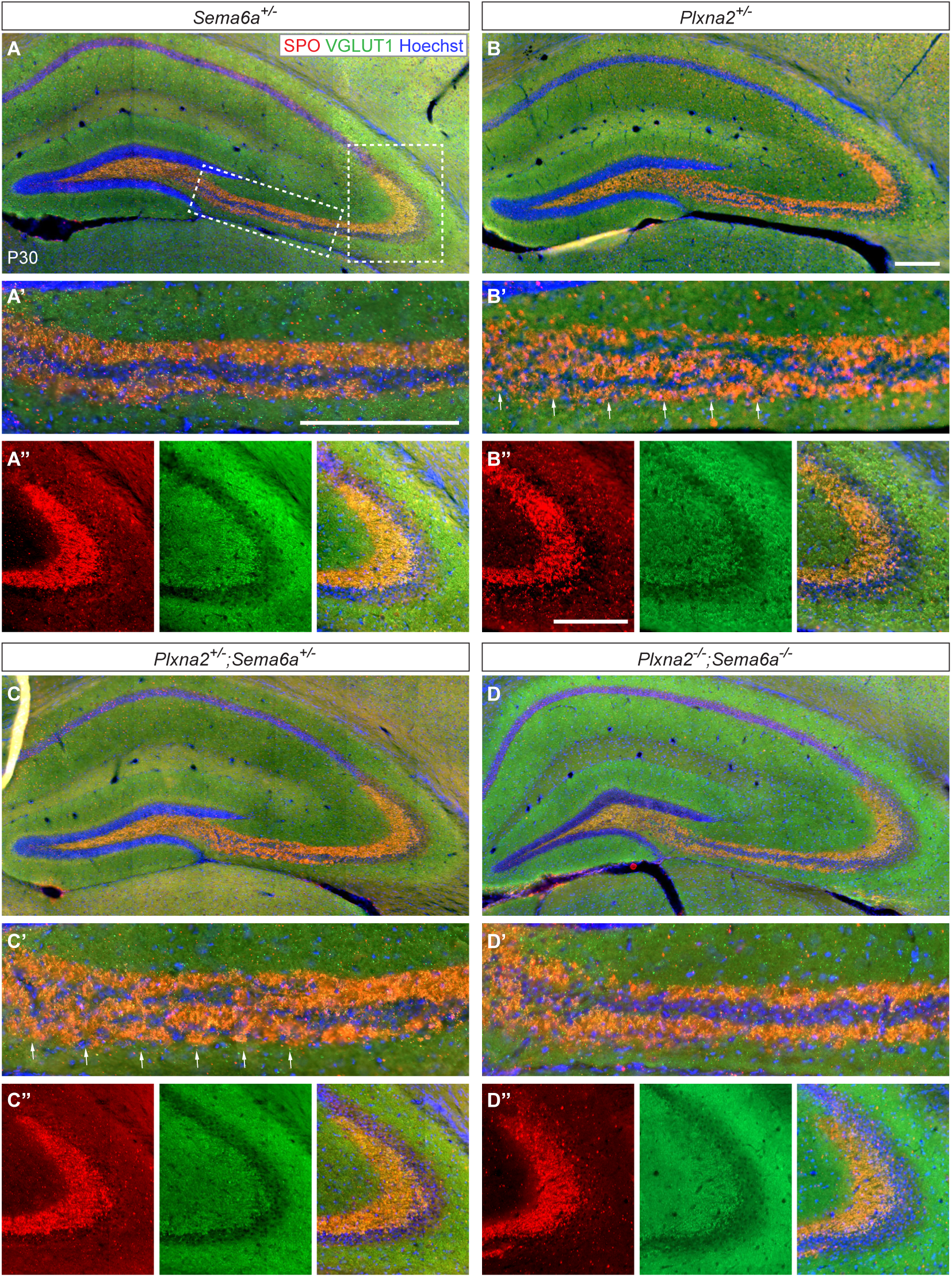
Loss of *Plxna2* results in *Sema6a* dependent and independent MF defects. (**A-D’’**) Representative images of coronal sections through the dorsal hippocampus of P30 (A) *Sema6a^+/−^* (n=4), (B) *Plxna2^+/−^* (n=5), (C) *Plxna2^+/−^*;*Sema6a^+/−^* (n=6), and (D) *Plxna2^−/−^*;*Sema6a^−/−^* (n=4) mice, stained with anti-SPO (red), anti-VGLUT1 (green), and Hoechst nuclear dye (blue). (**A’-D’**) Higher magnification images of area CA3c/b. Arrows point to region of defective MF bundling and positioning in *Plxna2^+/−^* and *Plxna2^+/−^*;*Sema6a^+/−^* mice, respectively. (**A’’-D’’**) Higher magnification images of area CA3a. Scale bar for all images, 200 µm. Quantification of MF defects is shown in Figure 5D and 5E.

**Figure 5.**
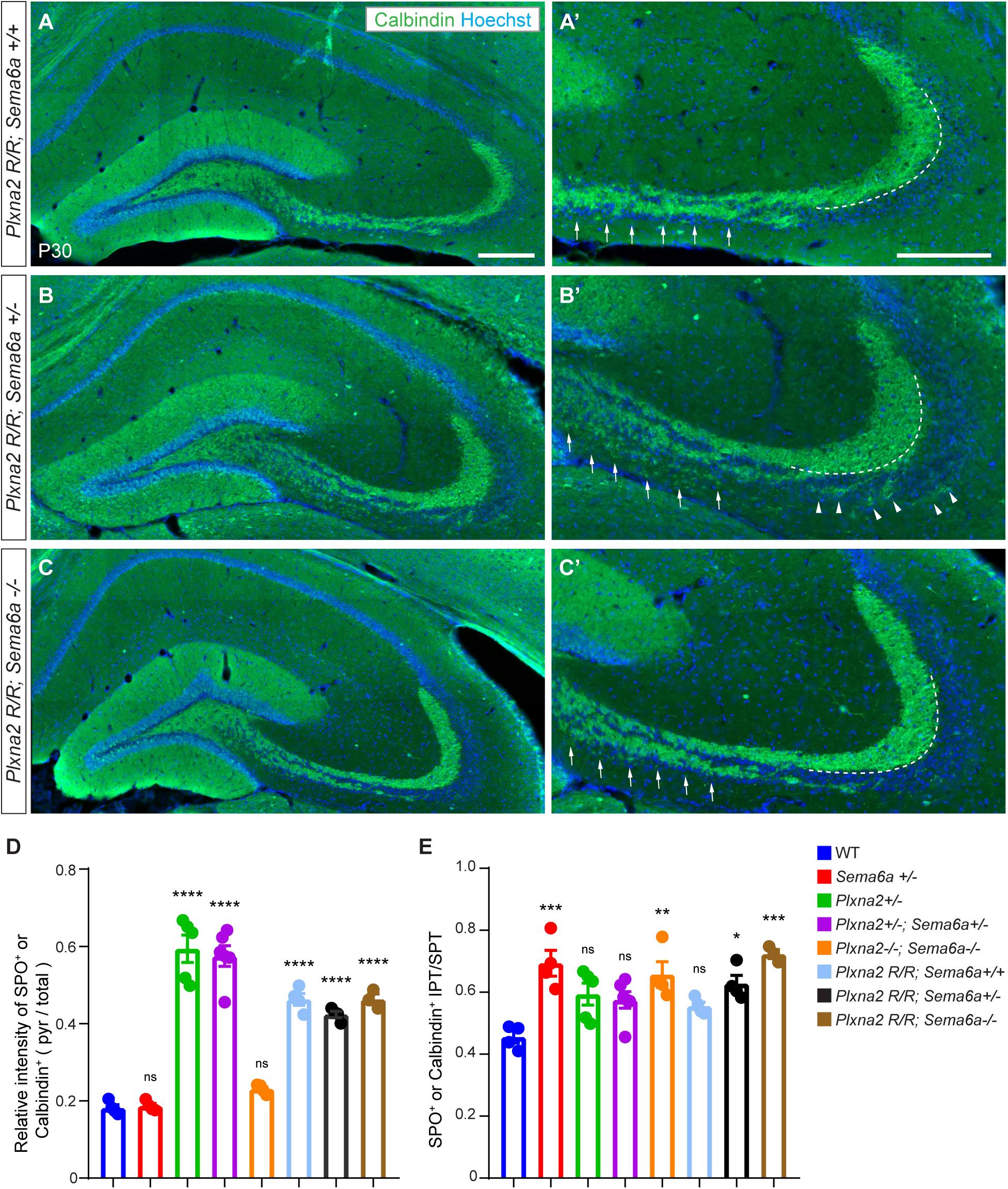
PlxnA2 GAP deficiency fails to rescue fasciculation defect caused by *Sema6a* knockout. (**A-C’**) Representative images of coronal sections through the dorsal hippocampus of P30 (A) *Plxna2^R/R^; Sema6a^+/+^* (n= 4), (B) *Plxna2^R/R^;Sema6a^+/−^* (n= 4) and (C) *Plxna2^R/R^;Sema6a^−/−^* (n= 3) mice, stained with anti-calbindin and Hoechst nuclear dye. (A’-C’) Higher magnification images of area CA3 shown in A-C. (A’) Arrows indicate hyper-fasciculation of MF in CA3c/b. (B’) Arrows in CA3c/b indicate aberrant MF defasciculation and arrowheads point to overextended IPT axons in CA3a (C’) Arrows in CA3c/b indicate area of incomplete MF partitioning. Scale bar, 200 µm. (**D**) Quantification of defective MF partitioning in area CA3c/b. Shown is the ratio of calbindin^+^ MFs in the pyramidal cell layer/total. (**E**) Quantification of aberrant IPT length. The ratio of IPT/SPT length is shown. Error bars are SEM. * p<0.05, ** p<0.01, *** p<0.001 and **** p<0.0001, ns, not significantly, one-way ANOVA.

Defects in MF partitioning are less pronounced in *Plxna2^R/R^* than in *Plxna2^−/−^* mice, indicating that PlxnA2 GAP-dependent and GAP-independent signaling events regulate specific aspects of MF patterning (**Figure 5A**). To explore whether MF defects in *Plxna2^R/R^* mice are sensitive to *Sema6a* gene dosage, we crossed *Sema6a* mice with *Plxna2^R^* mice. In *Plxna2^R/R^;Sema6a^+/+^* mice, the SPT and IPT fail to separate properly, and MF are ectopically positioned within the SP (**Figure 5A-A’**). Consistent with our previous findings (*22*), SPT axons in *Plxna2^R/R^* mice properly innervate SL in CA3a (**Figure 5A-A’**). Similar to *Plxna2^R/R^;Sema6a^+/+^* mice, the partitioning of the SPT and IPT is defective in *Plxna2^R/R^;Sema6a^+/−^* mice. However, in *Plxna2^R/R^;Sema6a^+/−^* mice the IPT is extended compared to *Plxna2^R/R^;Sema6a^+/+^* mice (**Figure 5A-B’**). In *Plxna2^R/R^;Sema6a^−/−^* mice, there is partial rescue of the SPT and IPT separation, compared to *Plxna2^R/R^;Sema6a^+/+^*, however IPT axons remain aberrantly positioned within the SP and IPT axons remain elongated in the SO (**Figure 5C-C’**). Together, these findings indicate that only some of the MF defects observed in *Plxna2^R/R^* mice are sensitive to *Sema6a* gene dosage (**Figure 5D-E**).

### Probing the molecular composition of the PlxnA2 receptor complex

The excessive fasciculation of MF axons in CA3 of *Plxna2−/−* mice suggests unmasking of an axon-axon adhesive function, normally counteracted by PlxnA2. Because similar defects were not observed in *Sema6a−/−* mice, this prompted further investigations into the PlxnA2 receptor complex. Specifically, we employed immunoproximity biotinylation *in vitro* on live, unfixed neurons to identify cell surface proteins in close proximity to, and potentially interacting with, PlxnA2 (*48, 49*). Primary neuronal cultures were prepared from whole hippocampus of neonatal WT and *Plxna2^−/−^* pups. Cultures were then incubated with antibodies directed against PlxnA2, and bound antibody was detected using an HRP-conjugated secondary antibody; *Plxna2^−/−^ hippocampal neurons showed no immunoreactivity using anti-PlxnA2 antibodies* (**Figure 6A**). Proteins in close proximity to PlxnA2 were then biotinylated by the addition of biotin-tyramide to the cultures. The PlxnA2 localized HRP generates the reactive biotin phenoxyl from biotin tyramide, resulting in the biotinylation of tyrosine residues in proximity to the HRP and PlxnA2, with a range of several hundred nm. Subsequently, cells were lysed, subjected to streptavidin pull-down, and analyzed by tandem-mass spectrometry (**Figure S6A**). PlxnA2 was identified as top hit in the anti-PlxnA2 dataset generated from WT cultures, and as expected, it was absent in the dataset obtained from *Plxna2^−/−^* cultures. Analysis of differentially abundant proteins revealed candidate components of the PlxnA2 receptor complex. Prominently featured were members of the Ig-CAM family such as neural cell adhesion molecule (NCAM), Neogenin-1 (Neo1), CXADR Ig-like cell adhesion molecule (CXARD), Contactin-1 (CNTN1), deleted in colorectal cancer (DCC), and L1-CAM (**Figure 6B**). The number of peptide spectral masses (PSMs) recovered for any given protein corresponds to 1) its amount and local density, 2) the number of extracellular tyrosines that can be biotinylated, and 3) its proximity to the biotinylation source. Importantly, the number of peptide spectral matches recovered showed no correlation to the number of extracellular tyrosines (**Figure 6B**), suggesting a stronger dependence of proximity and abundance on the number of recovered peptides. To estimate proximity to the biotinylation source, we calculated the ratio of extracellular tyrosines to PSMs for each identified protein. A lower ratio suggests a greater abundance of protein and closer proximity to the biotinylation source. We found that NCAM, CNTN1, and PlxnA4 were among those cell surface proteins with the lowest ratio (**Figure 6C**). Among the proteins identified with low ratios, many are expressed in developing granule cells and located to MF axons (*50*).

**Figure 6.**
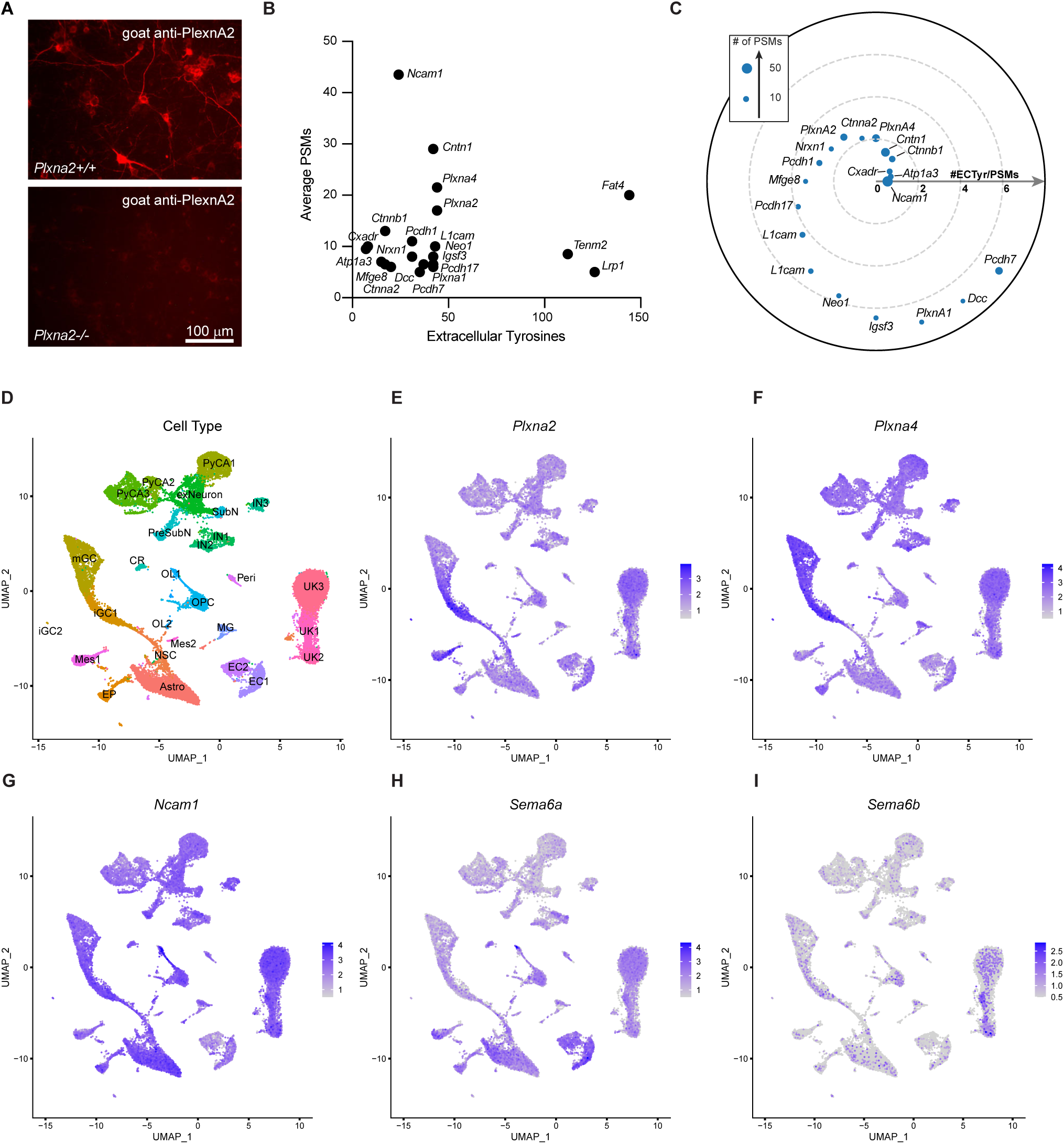
Cell surface interactome of PlxnA2 in primary hippocampal neurons. (**A**) DIV7 primary hippocampal neurons of *Plxna2^+/+^* and *Plxna2^−/−^* pups, incubated with anti-PlxnA2 antibody, followed by HRP mediated tyramine biotinylation of proximal surface proteins. Biotinylated proteins are stained in red. (**B**) Scatter plot of average peptide spectral matches (PSMs) in WT over *Plxna2^−/−^* cultures for biotinylated proteins identified by mass spectrometry plotted against the number of tyrosine residues in each protein’s extracellular domain. (**C**) Proximity plot showing biotinylated proteins ordered by extracellular tyrosine/PSM ratio. The plot is an estimate of abundance and proximity to the HRP secondary antibody bound to the PlxnA2 primary antibody. Each protein is represented by a circle with size proportional to the number of PSMs identified for that protein. (**D**) UMAP plots of integrated P7 and P10 mouse hippocampus snRNAseq datasets. Cell cluster labels are based on the presence of maker genes. (**E-I**) Feature plots for *Plxna2* (E), *Plxna4* (F), *Ncam1* (G), *Sema6a* (H) and *Sema6b* (I). Color coded calibration of relative gene expression is shown. Abbreviations; NSC, (neural stem cells), iGC (immature granule cells), mGC (mature granule cells), Astro (astrocytes/radial glia), PyCA (Pyramidal neurons in CA1, CA2, and CA3), SubN (subicular neurons), PreSubN (presubicular neurons), CR (Cajal retzius cells), IN (interneurons), Peri (pericytes), OPC (oligodendrocyte progenitors) OL (oligodendrocytes), EC (endothelial cells), MG (microglia), Mes (fibroblast-like cells), and UK (unknown).

For a detailed analysis of gene expression in the postnatal hippocampus, we took advantage of an existing snRNAseq dataset of mouse P10 hippocampus (*51*) and, in addition, carried out our own snRNAseq transcriptomic profiling of mouse P7 hippocampus. Datasets were subjected to dimensional reduction through principal component analysis. The first 20 principal components were used for shared nearest neighbor and Louvain cluster determination with a resolution parameter set at 0.5; uniform manifold approximation and projection analysis was used for cell cluster visualization and marker gene express was used for cell type identification (*29*) (**Figure S7**). Gene expression analysis revealed strong and preferential expression of *Plxna2* in immature granule cells (iGC), while *Plxna4* was abundant in iGCs and mature GCs (mGCs). *Ncam1* was strongly expressed by iGCs and mGCs, and its distribution was broader including additional cell types such as CA1-CA3 pyramidal neurons, interneurons and cells in the OL lineage (**Figure 6D-G**). *Sema6a* and *Sema6b* transcripts were detected in iGCs and mGCs, as well as CA1-CA3 pyramidal neurons, though at lower levels (**Figure 6D, H, I**). Dot plot analysis of additional candiate gene products identified by BAR shows their cell type specific distribution in the postnatal hippocampus (**Figure S7**).

Next, we selected several candiate molecules identified by the proximity biotinylation, including NCAM140, NCAM180, L1CAM, CHL1, Neogenin-1, DCC, CNTN1, and PCDH17, for further analysis. Using transient transfection of expression plasmids in HEK293T cells, we tested the ability of these transmembrane proteins to support Sema6A-Fc binding (**Figure S6E**) or to influence binding when co-expressed with PlxnA2. None of the candidates on their own supported binding of bath applied Sema6A-Fc, but neither did they abolish binding of Sema6A-Fc to PlxnA2. Because our estimate of protein proximity implied NCAM is the closest cell surface protein to the biotinylation source (**Figure 6C**), and since NCAM is expressed by GCs, localized to MFs, and is known to function in SPT and IPT separation (*52–54*), we decided to further explore this interaction.

### Genetic interactions between *Plxna2* and *Ncam1* in developing MFs

Though NCAM and PlxnA2 are broadly expressed by developing hippocampal neurons, co-immunoprecipitation (IP) studies failed to demonstrate a physical interaction between these two proteins (**Figure S6D**). This suggests the interaction is either too weak to be detected by IP or that it may occur indirectly, consistent with a 200-300 nm radius of biotinylated proteins relative to the antibody target (*49*). To further explore the functional relationship between NCAM and PlxnA2, we tested whether *Plxna2* and *Ncam1* interact genetically in developing neurons. It was shown previously that *Ncam1^−/−^* deficiency causes defects in SPT and IPT partitioning and IPT overextension (*52, 53*). We find that conditional *Ncam1* ablation in developing forebrain neurons, as assessed in *Ncam1^flox/flox^;Emx1-cre* mice, is sufficient to mimic these defects (**Figure 7A-B’**). MF partitioning is incomplete in *Ncam1^flox/flox^;Emx1-cre* mice, with axons ectopically positioned within the CA3c/b SP and few IPT axons that are overextended (**Figure 7B-B’**). No defects in MF partitioning were observed in *Ncam1^flox/flox^* processed in parallel (**Figure 7A-A’**) or in *Ncam1^flox/+^;Emx1-cre* mice, indicating that neuronal *Ncam1* is haplosufficient with respect to MF partitioning (**Figure 7A-C’**). However, there was a modest increase in IPT length in *Ncam1^flox/+^;Emx1-cre* mice. Similar to *Ncam1^flox/+^;Emx1-cre* mice, conditional ablation of one allele of neuronal *Plxna2* (*Plxna2^flox/+^;Emx1-cre*) results in normal MF patterning and a modest increase in IPT length (**Figure 7D-D’**). To ask whether *Ncam1* and *Plxna2* genetically interact, we generated compound heterozygous mice, *Plxna2^flox/+^;Ncam1^flox/+^;Emx1-cre* (**Figure 7E, E’**). Analysis of MFs revealed defective partitioning into the SPT and IPT bundles in CA3c/b, with many axons ectopically positioned within the SP. Laminar targeting of the SPT in CA3a appears normal, while the IPT is overextended (**Figure 7E-G**). Similar experiments with *Plxna4^+/−^* and *Ncam1^flox/+^;Emx1-cre* compound heterozygous mice (**Figure S8**) or *Sema6a^+/−^* and *Ncam1^flox/+^;Emx1-cre* compound heterozygous mice (**Figure S9**), failed to reveal genetic interactions. Taken together, these data show that neuronal *Ncam1* and *Plxna2* genetically interact and collaborate to regulate partitioning of SPT and IPT in CA3c as well as the length of the IPT.

**Figure 7.**
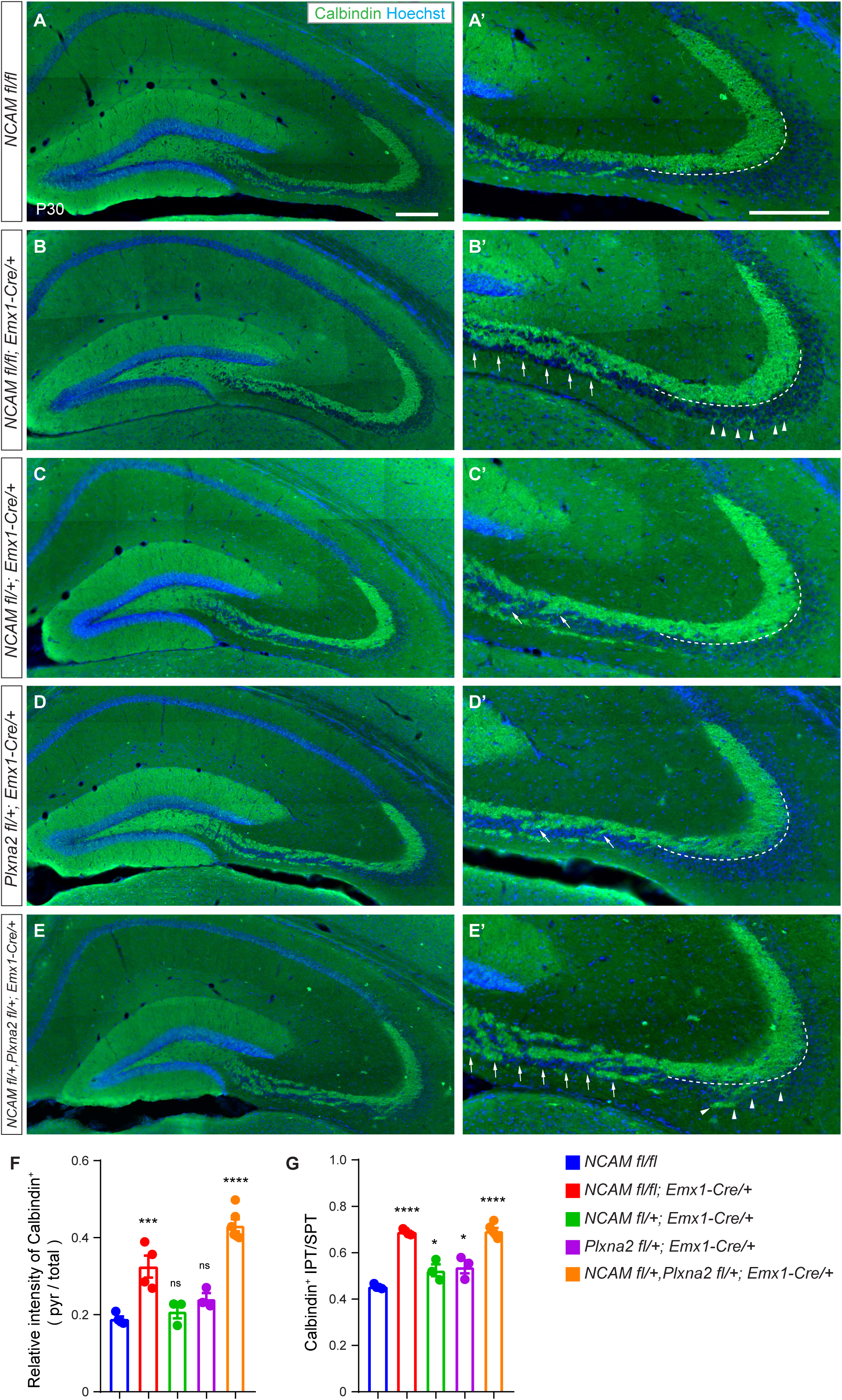
Genetic interaction of *Plxna2* and *Ncam1* in developing MFs. (**A-E**) Representative images of coronal brain sections through the dorsal hippocampus of P30 (A) *NCAM^fl/fl^* (n=4), (B) *NCAM^fl/fl^; Emx1-Cre/+* (n=4), (C) *NCAM^fl/+^; Emx1-Cre/+* (n=3), (D) *Plxna2^fl/+^; Emx1-Cre/+* (n=3) and (D) *NCAM^fl/+^, Plxna2^fl/+^; Emx1-Cre/+* (n=6) mice, stained with anti-calbindin and Hoechst nuclear dye. (**A’-E’**) Higher magnification images of area CA3 shown in A-E. (B’, C’, D’ and E’) Arrows in CA3c/b indicate regions with defective MF partitioning. (B’ and E’) Arrowheads in CA3a indicate overextension of IPT axons. Scale bar, 200 µm. (**F**) Quantification of defective MF partitioning in area CA3c/b. Shown is the ratio of calbindin^+^ MFs in the pyramidal cell layer/total. (**G**) Quantification of aberrant IPT length. The ratio of IPT/SPT length is shown. Error bars are SEM. * p<0.05, *** p<0.001 **** p<0.0001, ns, not significantly, one-way ANOVA.

## Discussion

We provide here novel insights into the molecular basis of MF patterning in the developing mammalian hippocampus. *Sema6a* is required for select guidance events, including axon partitioning and bundling into the SPT and IPT, and also avoidance of the SP in CA3c/b. In CA3a, *Sema6a* appears to be particularly important for regulation of IPT length. We show that neuronal *Sema6a*, but not OL-lineage-derived *Sema6a*, is necessary for proper MF organization and demonstrate a requirement for Sema6A reverse signaling. In *Plxna4^−/−^* mice, MF axon projections are severely defasciculated, resulting in aberrant innervation of CA3c, defasciculation of the SPT, innervation of the SR in CA3a, and overextension of the IPT. Several of these defects are replicated in PlxnA4 GAP mutants, highlighting the importance of forward signaling through GAP catalytic activity. In *Plxna2^−/−^* mice, MF axons fail to separate into SPT and IPT at the entry of CA3 and remain tightly bundled, resulting in ectopic innervation of the SP along the CA3c-CA3a axis. While axon bundling defects in CA3c persist in PlxnA2 GAP mutants, laminar targeting of the SPT to the SL in CA3a and length of the IPT appear largely normal. This shows that both PlxnA2 and PlxnA4 operate through GAP-dependent and GAP-independent mechanisms at specific sites along the MF trajectory. Analysis of the PlxnA2 proximity proteome in primary hippocampal neurons resulted in the identification of several Ig-CAM family members, and our analyses reveal that neuronal *Ncam1* and *Plxna2* genetically interact to regulate partitioning of MF axons into the SPT and IPT within CA3c/b. Neuronal *Plxna2* and *Ncam1* also interact to regulate IPT length, but not laminar targeting of the SPT to the SL. Together, our studies underscore the multifaceted functions of PlxnA2 and PlxnA4 signaling for proper navigation of MF axons at specific choice points along their trajectories, and they reveal a novel association between *Plxna2* and *Ncam1* for proper development of the DG-CA3 fiber system.

### Refined model of MF patterning

We describe three distinct guidance events that operate in a context-dependent manner and rely on specific aspects of PlxnA2 and PlxnA4 signaling (**Figure S10**). First, they act in the partitioning of MF projections into the SPT and IPT at the entry of CA3c and positioning of axon bundles immediately above and below the SP. Second, they regulate lamina-specific innervation of the SPT confined to the SL, and third, they control IPT length through stereotyped axon pruning. In *Plxna2^−/−^* mice, MFs are tightly fasciculated and fail to separate into SPT and IPT in CA3c, resulting in one prominent axon bundle ectopically positioned with the SP. In contrast in *Plxna4^−/−^* mice, MFs are severely defasciculated, failing to bundle into SPT and IPT at the entry of CA3c. This suggests that PlxnA4 signals “surround repulsion”, driving MF axon into fascicles, whereas PlxnA2 signals axon-axon repulsion important for MF partitioning (**Figure S10B**).

Loss of PlxnA2-GAP function is sufficient to mimic ectopic positioning of MFs within the SP, suggesting that PlxnA2 forward signaling through GAP catalytic activity is necessary for MF partitioning in CA3c. An observation at variance with the idea that Sema6A repulsive activity through PlxnA4 is competitively suppressed by binding to PlxnA2 (*11, 22*). PlxnA4-GAP activity is not required for avoidance of the SP but is necessary for MF axon bundling in the SPT and IPT. In CA3c, defects observed in *Plxna4^−/−^* are in part replicated in *Sema6a^−/−^* mice and following neuron-specific *Sema6a* ablation. This suggests that Sema6A/PlxnA4 forward signaling through GAP activation regulates axon fasciculation and separation into SPT and IPT, and that PlxnA4 GAP- independent mechanisms control avoidance of the SP. Some defects observed in *Plxna2^−/−^* mice are rescued in *Plxna2^−/−^;Sema6a^−/−^* double mutants, reminiscent of findings with *Plxna2^−/−^;Sema6a^lacZ/LacZ^* mice (*23*). Interestingly, only defects observed in CA3c/b of *Sema6a^−/−^* mice are rescued in *Plxna2^−/−^;Sema6a^−/−^* mice, while overextension of the IPT in *Sema6a^−/−^* is not sensitive to *Plxna2* deletion. This indicates an interdependence of *Sema6a* and *Plxna2* for MF patterning within CA3c/b, but not CA3a. Curiously, MF partitioning defects in CA3c of *Plxna2^R/R^* mice are not rescued in *Plxna2^R/R^*;*Sema6a^−/−^* mice, and the IPT is overextended, similar to *Sema6a^−/−^* mice. This suggests that selective disruption of PlxnA2 GAP catalytic activity results in dysregulation of PlxnA2-dependent signaling events that are independent of *Sema6a* gene dosage.

Longitudinal scRNAseq studies of late embryonic and early postnatal brains show preferential expression of *Sema6a* in interneurons and few excitatory neurons. This suggests that Sema6A produced within the SP, along the CA3c-CA3a axis, instructs MFs in a PlxnA4 dependent manner to separate into SPT and IPT bundles and avoid the SP. Our studies further reveal that PlxnA4 controls not only avoidance of the SP, but also promotes bundling of axons within the SPT and IPT along the CA3c-CA3a axis. Because avoidance of the SP is still observed in PlxnA4 GAP mutants, this suggests that Sema6A instructs MFs in a PlxnA4 GAP-independent manner. In CA3c of *Plxna4^−/−^* mice, expressivity of SPT axons defasciculating is more pronounced than in PlxnA4 GAP or *Sema6a^−/−^* mutants. This indicates that PlxnA4 regulates axon fasciculation within the SPT independently of Sema6A and independently of PlxnA4 GAP activity. Unlike the SPT, axons in the IPT require *Sema6a* function and PlxnA4 GAP activity for proper fasciculation, suggesting that PlxnA4 employs different signaling mechanisms to control axon fasciculation within the SPT and IPT. We speculate that the tight fasciculation of all MFs at the entry of CA3c observed in *Plxna2* mice is due to increased axon-axon adhesion, possibly involving multiple Ig-CAM family members as discussed below. The presence of PlxnA2 lacking GAP enzymatic activity is not sufficient to rescue the *Plxna2^−/−^* hyper-fasciculation phenotype in CA3c, indicating that PlxnA2 forward signaling through its GAP catalytic activity is necessary to overcome axon-axon adhesion (**Figure S10B**).

More distally, in area CA3a, SPT axons are strongly defasciculated in both *Plxna4^−/−^* and PlxnA4 GAP mutants, resulting in aberrant innervation of the SR. Because Sema6A and Sema6B have been shown to repel MF axons in a PlxnA4 dependent manner *in vitro* (*23*), and *Sema6a^−/−^* mice do not replicate the SPT defasciculation defects observed in *Plxna4^−/−^* mice, Sema6A and Sema6B likely collaborate to confine SPT axons to the SL *in vivo* (**Figure S10D**). Indeed, the number of aberrant MF projections is increased in *Sema6a^LacZ/LacZ^;Sema6b^−/−^* double mutants, compared to single mutants (*41*), yet less pronounced than in *Plxna4^−/−^* or PlxnA4 GAP mutants, possibly due to the hypomorphic nature of the *Sema6a^lacZ^* allele (*43*).

In early postnatal brains (P1-P7) Sema6A is abundant in the SLM, and thereby may direct SPT axons into the SL. In CA3a, absence of SPT axons from the SL and aberrant positioning within the SP of *Plxna2^−/−^* mice is no longer observed in PlxnA2 GAP deficient mice (*22*). These phenotypes are consistent with Sema6A being sequestered by PlxnA2, thereby mitigating Sema6A/PxnA4 repulsion (*11, 22*). In support, we find high levels of PlxnA2 immunoreactivity within the SL, but not SR or SP, at the time of SPT innervation. Thus, along the CA3c-CA3a axis, PlxnA2 may operate as a signaling receptor (CA3c/b) or as a competitive binding partner for Sema6A that attenuates PlxnA4 signaling (CA3a).

### Organizing principle for IPT stereotyped pruning?

Stereotyped pruning of the IPT has been studied extensively. The identification of Sema6A reverse signaling and PlxnA4 GAP activity as novel regulators of IPT length adds to a growing list of guidance molecules that shape MFs *in vivo* (**Figure S10C**). Defects in IPT guidance and laminar targeting were first reported in mice deficient for *Nrp2* (*55*), with subsequent studies revealing impaired stereotyped pruning of IPT axons (*56*). Pruning of IPT axons from temporary targets in CA3a is regulated by Sema3F, secreted from NPY^+^ interneurons in CA3, and requires IPT axons to express the canonical Sema3F receptors, Nrp2 and PlxnA3 (*39, 57*). Because many Semas function as growth inhibitory or repulsive axon guidance cues, Sema repulsive signaling was originally thought to break the distal most synaptic contacts of IPT axons thereby initiating stereotyped pruning. However, more recent evidence revealed that Sema3F/Nrp2/PlxnA repulsion-independent mechanisms underlie IPT pruning. Sema3F mediated IPT pruning requires the Rac-GAP protein β2-Chimaerin (Chn2) (*44*). Chn2 binds to the cytoplasmic domain of Nrp2 and is activated in a Sema3F dependent manner. Activation of Chn2 is necessary for axon pruning but dispensable for Sema3F mediated axon repulsion. In a similar vein, conditional ablation of the E3 ubiquitin ligase Cullin-5/RBX2 (CRL5) in the mouse forebrain results in IPT overextension at P21 and IPT defects persist into adulthood (*58*). Axons of primary GCs deficient for *Crl5* are strongly repelled by Sema3F, comparable to parallel processed WT GCs (*58*). Together these studies indicate that Sema3F mediated axon repulsion and stereotyped pruning can be dissociated at the molecular level.

Overextension of the IPT is observed in *Sema6a^−/−^* mice is partially replicated upon neuron specific, but not OL-lineage specific, *Sema6a* depletion. The less pronounced IPT defects in conditional mutants may reflect incomplete gene ablation or contributions from multiple cell types. Surprisingly, in *Sema6aΔcyt* mice the IPT is also extended, suggesting that Sema6A reverse signaling is involved. In a similar vein, CRMP family members (*59, 60*), and ephrin/Eph interactions are important not only for axon guidance but also for IPT pruning (*61, 62*). Ephrin-B reverse signaling through Grb4 and Rac was shown to regulate IPT pruning, with EphB family members functioning as redundant ligands for ephrin-B (*62*). The growing number of guidance cues and receptors that control IPT length is quite remarkable and questions whether they all are directly involved in stereotyped pruning or whether a more global mechanism is perturbed. For example, defects in axon guidance that result in aberrant synaptic contacts and altered circuit assembly may result in aberrant inhibitory or excitatory synapse numbers, changes in electrical activity, and altered network activity may ultimately impact IPT pruning (*63, 64*). Because Sema/Npn/Plxn and ephrin/Eph are known to regulate synaptic strength in the more mature nervous system (*65, 66*) changes in network activity may impair stereotyped pruning. IPT axons form collaterals to transiently synapse on basal dendrites of CA3 pyramidal cells during early postnatal development. Prior to IPT stereotyped pruning, these collaterals are removed in a *Plxna3*-dependent manner (*67*). A recent study found that neonatal exposure to ketamine results in extended IPT axons in juvenile mice. In these mice, unpruned synapses remained functional and exhibited increased spontaneous excitatory synaptic transmission, suggesting that elevated network activity may suppress IPT pruning (*68*). Changes in neuronal firing of dentate granule cells may be caused by altered input to dentate granule cells or reduced inhibition by interneurons. Additional studies are needed to determine if altered network activity in mice contributes to impaired IPT pruning. It is interesting to note that changes in network activity, such as are observed in epilepsy, have been shown to result in aberrant MF growth.

### NCAM1/Ig-CAMs and MF partitioning into SPT and IPT

There is evidence for extensive collaboration between Ig-CAM family members and Sema receptors for regulating neuronal morphology. Examples include *cis* and *trans* associations between L1-CAM and neuropilin-1 to control Sema3A responsiveness (*69*) and TAG1/CNTN2 regulated endocytosis of the Sema3A receptor complex (*70*). Subsequent studies identified CHL1 and Alcam as regulators of Sema signaling in midbrain dopamine neurons (*71*). NrCAM contributes to Sema3F-dependent dendritic spine remodeling (*72*) and CHL1 and Nrp2 function in Sema3B mediated dendritic spine pruning (*73*).

Using immunoproximity biotinylation with anti-PlxnA2 to find PlxnA2-associated proteins, we identified several Ig-CAM family members previously implicated in Sema signaling as well as new members that may be part of a larger receptor complex. The radius of biotinylation when using HRP is approximately 250 nm (*49*), and thus, surface molecules that do not directly associate with PlxnA2 may be detected. Binding studies in HEK293 cells revealed that none of the top candiate molecules identified supports direct binding of Sema6A-Fc. Gene expression analysis and work to date revealed prominent expression along developing MFs (*50*). *Ncam1* deletion in the mouse interferes with partitioning of SPT and IPT axons, resembling defects observed in *Plxna2^−/−^* and *Plxna2^R/R^* mice. Of interest, mice deficient for the intracellular monooxygenase MICAL-1, a known interaction partner of Plxn family members (*74, 75*), exhibited defects in MF partitioning and bundling in CA3c/b (*54*). Importantly, MICAL regulates NCAM function indirectly, by controlling growth cone targeting and surface expression of polysialic acid (PSA)-NCAM. Our previous studies showed that MICAL1 redox activity controls F-actin dependent surface trafficking of Rab6-positive vesicles laden with multiple Ig-CAMs (*54*). When coupled with the known associations between Plxn and MICAL family members and the genetic interaction between *Ncam1* and *Plxna2* we observe here, this suggests that the combined action of Sema6/PlxnA2 repulsion and NCAM mediated adhesion are necessary for MF partitioning and subsequent bundling of axons into the SPT and IPT. Our findings add to mounting evidence that Ig-CAM family member mediated axon-axon adhesive interactions are kept in check by Sema/Plxn mediated repulsion (*20, 21*). Of interest, proximity biotinylation with anti-PlxnA2 identified PlxnA4, however genetic studies did not find an interaction between *Ncam1* and *Plxna4*, indicating that only select PlxnA family members collaborate with *Ncam1* in developing neurons.

### Link to neuropsychiatric illness?

A growing number of mutations in *SEMA* and *PLXN* gene family members has been identified and associated with variable vulnerability to neuropsychiatric illnesses. This includes mutations in *PLXNA2* (*2, 4, 76*) and some of its ligands (*77–80*). In a similar vein, mutations in *NCAM1* (*81–85*) have been linked to neuropsychiatric disorders. Here we provide evidence that *Plxna2* and *Ncam1* collaborate to ensure proper brain wiring, establishing a new link between two powerful guidance systems. Our studies also highlight the intricacies and versatility of PlxnA2 and PlxnA4 mediated guidance decisions along the axonal trajectory of an individual neuron. *In vivo*, these include PlxnA forward signaling through engagement of GAP catalytic activity, as well as GAP independent guidance decisions, reverse signaling through Sema6A, and collaboration of PlxnA2 with Ig-CAM family members. Intrigulingly, many of these guidance events do not operate continuously or collaboratively but rather seem to function sequentially to ensure proper circuit wiring at specific choice points.

## Acknowledgements

We thank members of the Giger lab for critical reading of the manuscript and Isabel Brandt for excellent technical support. We thank Patricia Maness for *Ncam1 floxed* mice, Alex Kolodkin for *Sema6a floxed* mice, and Jean-Francois Cloutier for Ig-CAM expression plasmids. This work was supported by the National Institutes of Health R01MH119346 (RG), the NWO Gravitation program BRAINSCAPES: Roadmap from Neurogenetics to Neurobiology (NWO: 024.004.012) (RJP), and the Dr. Miriam and Sheldon G Adelson Medical Research Foundation (MR, AB, and RG).

## Material and Methods

### Animals

All procedures involving mice were approved by the University of Michigan Institutional Animal Care and Use Committee and performed in accordance with guidelines developed by the National Institutes of Health. *Plxna2^−/−^*, *Plxna2^flox/flox^*, the point mutant *Plxna2(R1746A); Plxna2^R^* allele, and *NCAM^flox/flox^* mice have been described (*13, 22, 86, 87*). *Sema6a ^flox/flox^* mice (*43*), *Sema6aΔcyt* (*40*). The point mutant *Plxna4(R1745A); Plxna4^R^* allele, was generated by CRISPR/Cas9 mediated gene editing, using established methods (*22*). For additional information see Supplemental Material.

### Histological Procedures

Animal perfusion, tissue preparation and staining were carried out as described (*22*). Mouse pups at postnatal day (P)1, P7 and P30 were collected. Length of MF projections, and distribution of SPO^+^ or Calbindin^+^ MFs within CA3 of the hippocampus was assessed using ImageJ.

### Ligand-receptor binding studies

Binding assay was done as described previously (*13, 22*)

### Immunoproximity biotinylation and Mass Spectrometry

Immunoproximity biotinylation was performed as described previously (*49*).

### snRNA-sequencing

Single-nuclear RNA sequencing and data analysis were carried out as described previously (*88, 89*).

### Statistical Analysis

GraphPad Prism 8 was used for data analysis. A two-tailed unpaired Student’s *t* test was used for single comparison. One-way ANOVAs were performed for multiple comparisons. Details on statistical analyses are provided in figure legends. p<0.05 was considered statistically significant. Details on “n” representation and exact value are included in figure legends.

