## Supplemental Information for "Diverse and Location-Specific Roles of PlexinA2, PlexinA4, and NCAM in Developing Hippocampal Mossy Fibers"

Supplemental Information includes a detailed description of experimental procedures and ten figures.

#### *Plxna4(R1745A)* mice; Plxna4R allele

The point mutation was introduced into the mouse *Plxna4* locus by CRISPR/Cas9-mediated gene editing. Briefly, single strand DNA fragments (sgDNA, 5'- CTCTATACAGCCTGCCCCTACGG -3') and ssODN template (5'- ACCCAACCACTCCTGACGAAAGCTGCTTCTTTCCCTCTATACAG**CCTGCCaCTtgccTTcTGGGTGAATATG ATCAAGAACCCTCAGTTTGTGTTTGACATCCATAAGAACA** -3') containing the desired mutations (CGG→GCC) were synthesized and purified by Integrated DNA Technologies. *In vitro* transcription (IVT) for sgRNA was performed with MEGAshortscript T7 kit (Life Technologies) using sgRNA template cloned in px330 plasmid (Addgene plasmid 42230); the IVT product was purified using MEGAclear kit (Life Technologies). A mixture of Cas9 mRNA (TriLink Biotechnologies, 100 ng/µl), sgRNA (50 ng/µl), and ssDNA (100 ng/µl) was injected into fertilized eggs from C57BL/6J mice (Jackson Laboratory, Stock #000664). Viable two-cell stage embryos were transferred to pseudo-pregnant ICR females to generate six founder mice which were subsequently bred with C57BL/6 mice for germline transmission to generate F1 mice. Five correctly targeted F1 mice were identified and two of these founders were analyzed and maintained on a C57BL/6 background. Mice carrying the R1745A point-mutation in Plxna4 were identified by PCR and independently confirmed by DNA sequencing. For PCR genotyping the following primers were used: Forward, 5'- GGTCCCAGCAACAGGCAGATTGGA -3'; Reverse-WT, 5'- GGGTTCTTGATCATATTCACCCAAAACC -3'; Reverse-Mutant, 5'- GGGTTCTTGATCATATTCACCCAgAAggca -3' and amplification conditions: 94°C for 30 seconds, 70°C for 45 seconds, 72°C for 1 minute, and repeat for 35 times.

#### HEK293 cell ligand-receptor binding assay

Ligand-receptor binding studies in HEK293 cells were carried out as described (*1*). Briefly, HEK293 cells, cultured in 24-well plates, were transiently transfected with the following expression constructs, *eGFP, Sema6a, Plxna2, Plxna4, Plxna4*$^{\Delta cyto}$*, Plxna4*$^{R}$*, Ncam140, Ncam180, Cntn1, L1CAM, Neo1, Dcc, Pcdh17* and *Chl1* using lipofectamine2000 (*1*). Thirty-six hours after transfection, cells were rinsed in OptiMEM and incubated for 75 min at room temperature with mouse Sema6A-Fc (R&D Systems, cat# 9017-S6-050) at a final concentration of 2 µg/ml in OptiMEM.  For detection of bound Sema6A-Fc, anti-human IgG1 conjugated to human alkaline phosphatase (AP) was used. The AP reaction was developed in the presence of NBT/BCIP substrate (Sigma) at pH 9.5 and stopped by rising with PBS.

**Primary neuronal cultures**

Embryonic-day (E) 17.5-18.5 mouse brains were dissected in Leibovitz's L-15 medium (Invitrogen 21083-027) and transferred to individual wells of a 12-well plate containing ice-cold Hibernate-E solution (Invitrogen A12476-01), 10% fetal bovine serum (Atlanta Biologicals) and 2% B27 supplement (Invitrogen 17504-044). For PCR genotyping, tail samples from each embryo were biopsied. While genotyping was performed, embryos were kept on ice for approximately 3.5 hours. Then, hippocampi of embryos with the same-genotype were pooled in Leibovitz's L-15 dissection medium. To remove $Ca^{2+}$ and $Mg^{2+}$ from solution, hippocampi were gently rinsed twice in Hank's Balanced Salt Solution (HBSS) (Invitrogen 14175-095) and digested for 15-20 min at 37°C in HBSS containing 0.25% Trypsin/EDTA (Invitrogen 15400-054) and 0.01% DNase I (Roche 10104159001) with gentle agitation. Trypsin was inhibited by incubation for 5 min in ice-cold Dulbecco's Modified Eagle Medium (DMEM) (Invitrogen 11960-044) containing 10% FBS. Samples were rinsed once in the same solution and transferred to ice-cold Neuronal Growth Medium [NGM: Neurobasal (Invitrogen 21103-049) with 25 mM D(+)-glucose, 2 mM Glutamax I (Invitrogen 35050-061), 50 units/mL penicillin and 50 µg/mL streptomycin (Invitrogen 15140-122), and 2% B27 supplement]. After another rinsing step in NGM, samples were centrifuged at 100 *x g* for 4 minutes and re-suspended in 1 mL of NGM. A fire-polished Pasteur pipette was used for trituration, 15 passes, to obtain a cell suspension. Cells were incubated with Trypan blue solution (Invitrogen 15250-061) and a hemacytometer was used for counting of live cells. Approximately 600,000 cells were plated per well of a 6-well plate (TPP TP92006), coated with of 100 µg/mL poly-D-lysine (Sigma P7886). Cells were culture in 2 mL of NGM per well. On day in vitro (DIV) 3, a half-media change with pre-warmed astrocyte-conditioned media (*2*), supplemented with 2% B27, was carried out.

**Immunoproximity biotinylation and Mass Spectrometry**

Experiments were carried out as described previously (*3*). At DIV7, primary hippocampal neurons prepared from WT and *Plxna2*$^{-/-}$ embryos, were incubated with anti-PlxnA2 antibody for 1 hr at 37°C. After a brief rinse in Neurobasal medium, neurons were incubated with HRP-conjugated secondary antibody for 30 minutes at 37°C. Following rinsing in ice-cold PBS, neurons were incubated with TSA working solution (PerkinElmer, NEL749B001KT) for 5 min, and the reaction was quenched with freshly made 0.5 M sodium ascorbate for 5 min. After rinsing with PBS, neurons were lysed with RIPA buffer, and lysates centrifuged at 12500 x g for 30 min at 4°C. Supernatants were incubated with streptavidin beads for 2 hrs and biotinylated proteins precipitated by centrifugation.  Pellets were rinsed and analyzed by Mass Spectrometry as described previously (*3*).

**Western blot analysis**

Brains were lysed in radioimmunoprecipitation assay (RIPA, Sigma R0278) buffer supplemented with 50 mM BGP, 1 mM $Na_3VO_4$, and 1:100 PIC. Brains were further homogenized using a pestle motor mixer (RPI 299200) and incubated on ice for 30 minutes with intermittent vortexing. After max spin in a refrigerated table-top centrifuge, supernatants were combined with equal volumes of 2x Laemmli sample buffer with BME, boiled for 15 minutes, and stored at -80°C. Samples, 5 µg per lane, were loaded and separated by SDS-PAGE and transferred onto PVDF membrane (Millipore IPVH00010). Membranes were incubated in 2% blotting-grade blocker (BioRad 1706404) prepared in 1x TBS-T (TBS pH 7.4, containing 0.1% Tween- 20) for 2 hours at room temperature, and probed overnight with the following primary antibodies diluted in 2% BSA (Fisher Scientific BP1600): α-PlxnA2 (R&D Systems AF5486, 1:1000), α-PlxnA4 (R&D Systems MAB5856, 1:1000), and α-βIII Tubulin (Promega G7121, 1:10000). Horseradish peroxide (HRP)-conjugated secondary IgG antibodies (EMD Millipore) were diluted in 2% BSA, and the HRP signal was developed using chemiluminescent substrates from Thermo Fisher Scientific (34080 or 34095) and Li-COR Biosciences (926-95000). Protein bands were visualized and quantified using LI-COR C-Digit and Image Studio software.

**Immunofluorescence labeling**

P1, P14, P30, and adult mice were perfused transcardially with ice-cold 4% paraformaldehyde (PFA) in phosphate buffered saline pH7.4 (PBS). Brains were extracted, post-fixed in perfusion solution for 2hrs, cryoprotected in 30% sucrose in PBS and frozen in OCT. Coronal sections were cut at 18 µm or 50 µm thickness and used for mounting on Superfrost microscope slides or free-floating staining, respectively. For immunofluorescence labeling, sections were incubated in 5% horse serum, 0.1% Triton X-100 in PBS for 30 minutes at room temperature followed by Incubation in primary antibody at 4ºC overnight. Primary antibodies were used at the following dilutions: goat anti-PlxnA2 (1:500, R&D Systems); mouse anti-PlxnA4 (1:500, R&D Systems); chicken anti-GFAP (1:1000, Aves Lab); rabbit anti-HA (1:500, Cell Signaling); mouse anti-MAG (1:500, Millipore); rat anti-MBP (1:500, Millipore); rat anti-PDGFRα (1:500, BD Pharmingen); rabbit anti-Calbindin (1:500, Swant); rabbit anti-Synaptoporin (SPO; 1:500, Synaptic Systems); guinea pig anti-VGLUT1 (1:500, Millipore). Species-specific and Alexa Fluor-conjugated secondary antibodies (Life Technologies) were used at 1:500. Nuclei were labeled by incubation in Hoechst 33342 (Invitrogen). Sections were mounted and imaged with a Zeiss Apotome2 microscope equipped with a Axiocam 503 mono camera and Zen 2 software.

**snRNA sequencing**

Single-nuclear RNA sequencing and data analysis were carried out as described previously (*4, 5*). Briefly, P7 WT mouse hippocampi were micro-dissected and tissue homogenized with a dounce homogenizer in ice-cold EZ PREP buffer (Nuclei isolation kit, Sigma # NUC101-1KT). Samples were briefly centrifuged, rinsed, and resuspended in buffer containing 0.1% RNase inhibitor (Roche, #03335399001). After filtering through a 40 µm cell strainer, the number of nuclei was counted using a hemocytometer. For barcoding and library preparation, we used the Chromium Next GEM Single Cell 3' Reagent kit v3.1 (Dual Index). Barcoding and library preparation was performed following the manufacturer's protocols. Briefly, to generate single-nuclei gel-bead-in-emulsion (GEMs) solution, approximately 15,000 nuclei, in a final volume of 43 µl, were loaded on a Next

GEM Chip G (10x Genomics) and processed with the 10x Genomics Chromium Controller. Reverse transcription was performed as follows: 53°C for 45 minutes and 85°C for 5 minutes in a Veriti Therml Cycler (Applied Biosystems). Next, first-strand cDNA was cleaned with DynaBeads MyOne SILANE (10x Genomics, 2000048). The amplified cDNA and final libraries were prepared and cleaned with SPRIselect Regent kit (Beckman Coulter, B23318). A small aliquot of each library was used for quality control to determine fragment size distribution and DNA concentration, using a bioanalyzer. Libraries were pooled for sequencing with a NovaSeq 6000 (Illumina) at an estimated depth of 50,000 reads per nucleus.

**snRNAseq data analysis**

Raw snRNAseq data sets were processed using the 10x Genomics CellRanger software version 7.0.1. The CellRanger “mkfastq” function was used for de-multiplexing and generating FASTQ files from raw BCL. The CellRanger “count” function with default settings was used with the mm10 reference genome supplied by 10x Genomics, to align reads and generate single nuclei feature counts. CellRanger filtered nuclei and counts were used for downstream analysis in Seurat version 4.0.5 implemented in R version 4.1.2. Nuclei were excluded if they had fewer than 500 or more than 7,500 unique features. The CellRanger output from snRNAseq dataset of P10 (GEO: GSE186216) (*9*) was downloaded and cells with unique features between 500 and 10,000 were kept. The public P10 and our P7 data were integrated and normalized following a standard Seurat SCTransform+CCA integration pipeline (*6*). Principal component analysis was performed on the top 3000 variable genes and the top 25 principal components were used for downstream analysis. A K-nearest neighbor graph was produced using Euclidean distances. The Louvain algorithm was used with resolution set to 0.5 to group cells together. Non-linear dimensional reduction was done using UMAP. The top 100 genes for each cluster, determined by Seurat’s FindAllMarkers function and the Wilcoxon Rank Sum test, were submitted to Qiagen’s Ingenuity Pathway Analysis (IPA) software – version 70750971 (Qiagen Inc., https://digitalinsights.qiagen.com/IPA) using core analysis of up- and down-regulated expressed genes. Top-scoring enriched pathways, functions, upstream regulators, and networks for these genes were identified utilizing the algorithms developed for Qiagen IPA software (*7*), based on Qiagen’s IPA database of differentially expressed genes.

**Statistical Analysis**

GraphPad Prism 8 was used for data analysis. A two-tailed unpaired Student’s *t* test was used for single comparison, and one-way ANOVA was used for multiple comparisons. $p<0.05$ was considered statistically significant.

### Supplementary figures

#### Figure S1. MF defects in *Sema6a-/-* mice persist through adulthood

(A) Representative image of coronal brain section at the level of the dorsal hippocampus of a 9-month-old WT mouse (n= 2). (B). Representative image of coronal brain section at the level of the dorsal hippocampus of a 9-month-old and *Sema6a-/-* mouse (n= 2). Nuclei are labeled in blue (Hoechst dye) and MF projections with anti-Calbindin (green). In (B), arrows point to the region where MF axons fail to properly partition into the SPT and IPT in CA3c/b. Arrowheads in CA3a indicate overextension of IPT axons. Scale bar, 200 µm.

#### Figure S2. Longitudinal analysis of *Plxna2, Plxna4, Sema6a,* and *Sema6b* expression

(A) UMAP plot of Integrated snRNAseq datasets generated from E10, E11, E12, E13, E14, E15, E16, E17, E18 (S1), E18 (S3), P1, and P4 mouse cerebral cortex. Color coding is used to show cell distribution at each developmental time point. Feature plots are shown for (B) *Rbfox3* (NeuN) post-mitotic neurons, (C) *nes* (Nestin) neural progenitor cells, (D) *Gad1* (GAD67) interneurons, (E) *Plxna2,* (F) *Plxna4,* (G) *Sema6a,* (H) and *Sema6b*. Calibrated gene expression levels are shown. Datasets were originally described in (*8*).

#### Figure S3. Complementary distribution of Sema6A and PlxnA2 in the early postnatal hippocampus

(A-D) Coronal sections through the P1 hippocampus of *Sema6a(fl/fl)* mice stained with anti-PlxnA2 and anti-HA (Sema6A). Hoechst dye (blue) was used to visualize nuclei. (D) Merged staining is shown and reveals largely complementary distribution of PlxnA2 (green) and Sema6A (magenta). (E-H) Higher magnification of the region in the white dotted square shown in D. The dotted line demarcates the boundary between the stratum pyramidale (Hoechst labeled nuclei) and the stratum lucidum (SL). (I-L) Coronal sections through the P7 hippocampus showing higher magnification images of area CA3. The dotted line demarcates the boundary between the stratum pyramidale (Hoechst labeled nuclei) and the SL. Scale bar, 200 µm (A-D); 20 µm (E-L).

#### Figure S4. Analysis of Sema6A distribution in glial cells

(A, B) Coronal brain sections of P7 *Sema6a(flox/flox)* mice, stained with anti-HA to visualize distribution of Sema6A. In the hippocampus, HA-Sema6A is most abundant in the stratum lacunosum-moleculare (SLM) and in the dorsal thalamus, the lateral geniculate nucleus (LGN). Hoechst dye (blue) was used to visualize nuclei. (C-F) Coronal sections through the P7 neocortex. Double immunofluorescence labeling with anti-HA (Sema6A), anti-MAG, and anti-MBP (mature myelin producing oligodendrocytes). (G-J) Double immunofluorescence labeling with anti-HA (Sema6A), anti-PDGFRα (oligodendrocyte progenitor cells), and anti-GFAP (astrocytes/radial glia). Scale bar, 200 µm in (A, B); 50 µm in (C-J).

#### Figure S5. PlxnA4(R1745A) is localized to the cell surface and supports Sema6A binding in *trans* and *cis* configuration

(A) Anti-PlxnA4 Western blot analysis of HEK293 cells transiently transfected with expression plasmids for green fluorescent protein *(eGfp), Plxna4, Plxna4(R1745A),* and *Plxna4* lacking the cytoplasmic domain

(*Plxna4Δcyto*). Anti-actin is shown as loading control. (B) Binding of Sema6A-Fc fusion protein to HEK293 cells transiently transfected to express recombinant PlxnA4, PlxnA4(R1745A), PlxnA2 and eGFP (top raw); and following co-expressing of full-length Sema6A (bottom raw). Scale bar, 50 μm. (C) Western blot analysis of forebrain lysates prepared from P7 *Plxna4*$^{R/R}$, *Plxna4*$^{R/+}$ and *Plxna4*$^{+/+}$ pups, probed with anti-PlxnA4. Anti-actin is shown as loading control.

**Figure S6. Workflow for immunoproximity biotinylation and Mass Spectrometry**
(A) Workflow for anti-PlxnA2 antibody proximity biotinylation in DIV7 hippocampal neurons. (B-D) Western blot analysis of biotinylated proteins labeled using anti-PlxnA2, anti-Goat-IgG-HRP (secondary only), or Sema5A-Fc. Streptavidin-HRP was used for detection of biotinylated proteins. (C) Western blot of PlxnA2 pull-down from DIV7 hippocampal neurons probed with anti-PlxnA2. (D) Anti-PlxnA2 IP of DIV7 hippocampal lysates probed with anti-NCAM. IP, immunoprecipitate; PIP, post immunoprecipitation supernatant. (E). Binding of Sema6A-Fc to HEK293 cells transiently transfected with expression constructs for *Plxna2* and candiate receptor components *NCAM140, NCAM180, Cntn1, L1CAM, Neo1, DCC, Pcdh17, and CHL1*. Scale bar, 200 μm.

**Figure S7. Analysis of guidance cue expression in the postnatal hippocampus.**
(A) Dotplot analysis of integrated snRNAseq datasets generated from P7 and P10 mouse forebrain showing calibrated expression of marker gene products used for cell type identification. (B) Dotplot analysis of gene expression of *Plxna1-4*, *Sema5a, Sema5b, Sema6a,* and *Sema6B* and candidate gene products identified by BAR, including *Ctnnb1 (catenin beta1), Tenm2 (teneurin transmembrane protein 2), Neo1 (neogenin 1), Cntn1 (contactin 1), Ctnna2 (catenin alpha2), Nrxn1 (neurexin 1), Igsf3 (immunoglobulin superfamily member 3), Dcc (DCC netrin 1 receptor), Mfge8 (milk fat globule EGF and factor V/VIII domain containing), Pcdh1 (protocadherin 1), L1cam (L1 cell adhesion molecule), Cxadr (CXADR Ig-like cell adhesion molecule), Lrp1 (LDL receptor related protein 1), Fat4 (FAT atypical cadherin 4), Pcdh7 (protocadherin 7), Atp1a3 (ATPase Na/K transporting subunit alpha 3), Pcdh17 (protocadherin 17)*. Gene expression levels are normalized to average gene expression (color coded calibration). For each cell cluster, the percentile of cells expressing a specific gene product is indicated by the dot size. The snRNAseq dataset of P10 (GEO: GSE186216) (*9*) was integrated with our P7 mouse dataset. Astrocytes (Astro), neural stem cells (NSC), ependymal cells (EP), immature granulate cells (iGC), mature granule cells (mGC), pyramidal neurons (Py), Cornu Ammonis (CA), excitatory neurons, non-pyramidal (exNeuron), interneurons (IN), Cajal-Retzius cells (CR), presubicular neurons (PS), subicular neurons (SN), oligodendrocyte progenitor cells (OPC), microglia (MG), endothelial cells (EC), pericytes (Peri), mesenchymal cells (Mes), unknown clusters (UK).

**Figure S8. Lack of evidence for genetic interaction between *Plxna4* and *NCAM*.**
(A-C') Coronal brain sections through the dorsal hippocampus of P30 (A) *PlxnA4+/-* (n=5), (B) *Ncam1 flox/+; Emx1-Cre/*+ (n=3), and (C) *PlxnA4+/-, Ncam1 flox/+; Emx1-Cre/*+ (n=4) mice, stained with anti-Calbindin. Scale bar, 200 μm. (D) Quantification of MF partitioning in CA3c/b. Shown is the ratio of anti-calbindin in

pyr/total. (E) Quantification of the ratio of IPT/SPT length. Error bars are SEM. ns, not significantly, one-way ANOVA.

**Figure S9. *Sema6a* does not genetically interact with *NCAM.***

(A-B') Coronal brain sections through the dorsal hippocampus of P30 (A) *Ncam1 flox/+, Sema6a flox/+* (n=3) and (B) *Ncam1 flox/+, Sema6A flox/+; Emx1-Cre/+* (n=4) mice, stained with anti-Calbindin to visualize MF. Scale bar, 200 µm. (C) Quantification of MF partitioning in CA3c/b. Shown is the ratio of anti-calbindin in pyr/total. (D) Quantification of the ratio of IPT/SPT length. Error bars are SEM. ns, not significantly, one-way ANOVA.

**Figure S10. Graphical summary of guidance events that orchestrate MF patterning**

(A) Overview of MF projections in the mature hippocampus of WT mice. The SPT shows laminar targeting to the SL along the CA3c-CA3a axis and the IPT shows laminar targeting to the SO within CA3c and CA3b. (B) As MF axons exit the hilus and begin to innervate area CA3s, they split into two prominent fascicles, the SPT and IPT. We show that **surround repulsion** mediated by Sema6A (and possibility Sema6B), through PlxnA4 forward signaling and GAP catalytic activity prevents SPT axons from innervation of the SR. Neuronal *Ncam1* is necessary for formation of distinct MF fascicules, that wander above (SPT) and below (IPT) the CA3 pyramidal cell layer (Pyr). We propose that homophilic NCAM-NCAM association promotes **axon-axon adhesion**. NCAM mediated axon bundling is countered by Sema6A-PlxnA2 forward signaling through GAP catalytic activity. (C) In CA3b, **stereotyped pruning** of IPT axons is impaired in *Sema6a-/-*, *Sema6aΔcyt*$^{\Delta/\Delta}$, *Plxna4-/-*, *Plxna4*$^{R/R}$ and *Ncam1* cKO mice. (D) Laminar targeting of SPT projections to the SL requires surround repulsion through Sema6A-PlxnA4-GAP forward signaling in MF axons. PlxnA2 produced by pyramidal neurons is enriched in the SL, and may **sequester** Sema6A, thereby attenuating Sema6A-PlxnA4 repulsion, allowing SPT axons to innervate the SL.

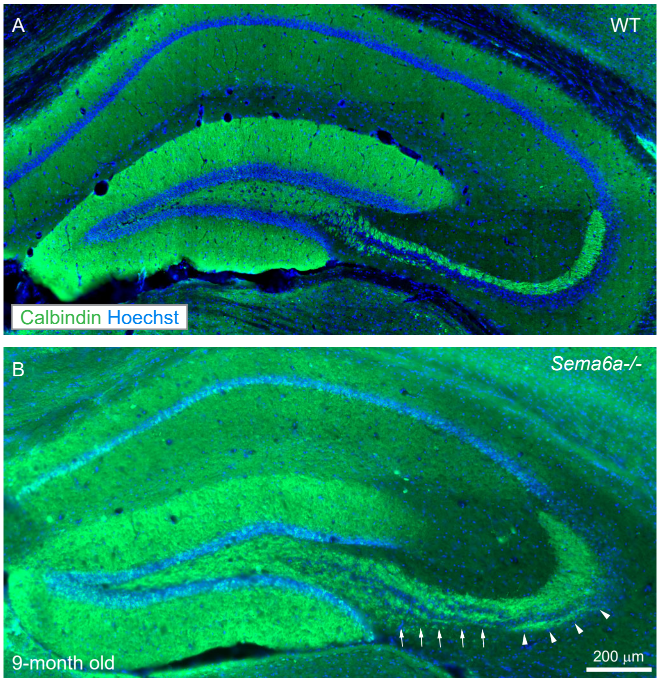


**Figure S1.**

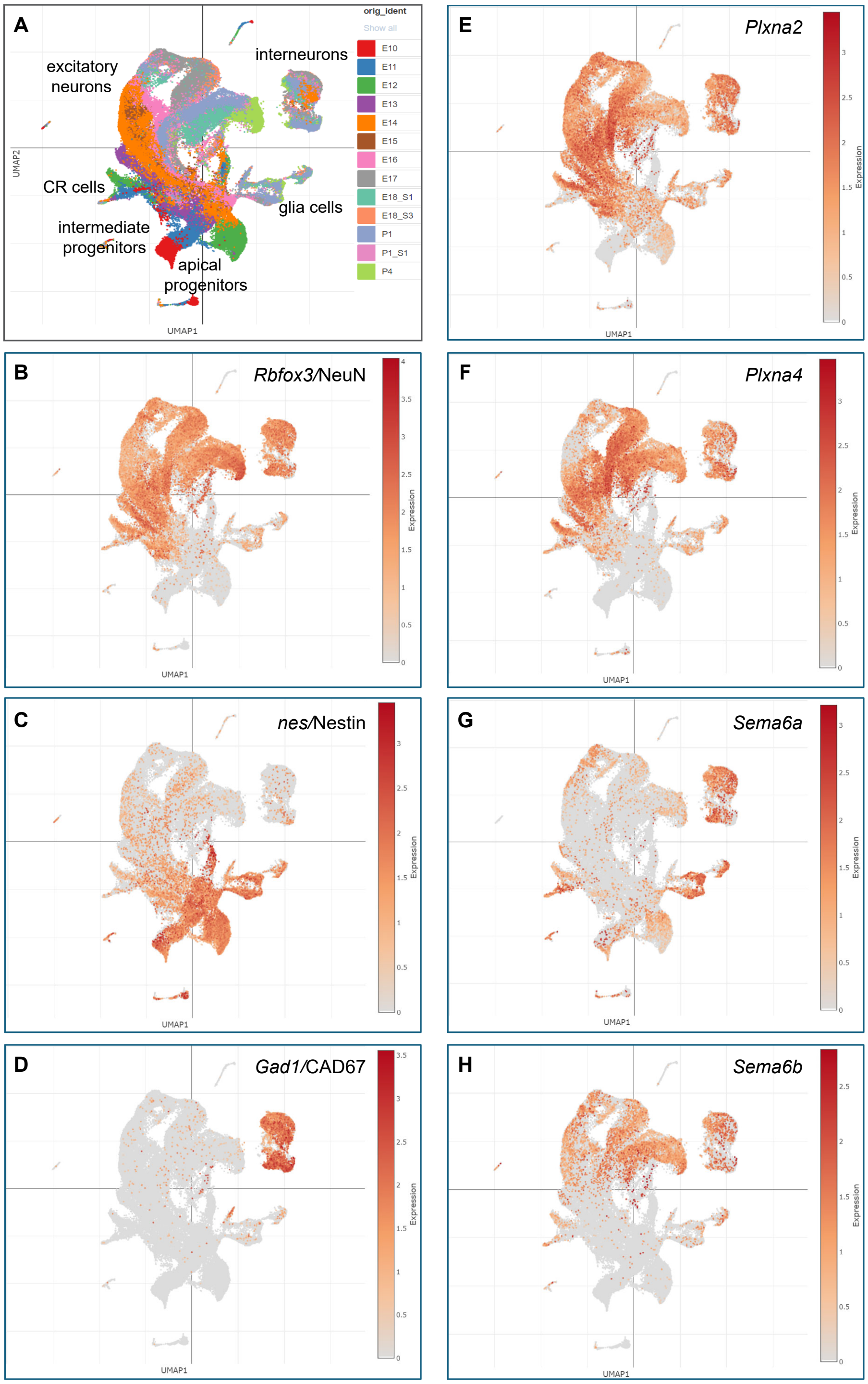
A
excitatory neurons
interneurons
CR cells
glia cells
intermediate progenitors
apical progenitors
orig_ident
Show all
E10
E11
E12
E13
E14
E15
E16
E17
E18_S1
E18_S3
P1
P1_S1
P4
UMAP1
UMAP2
B
Rbfox3/NeuN
C
nes/Nestin
D
Gad1/CAD67
E
Plxna2
F
Plxna4
G
Sema6a
H
Sema6b
Expression

**Figure S2.**

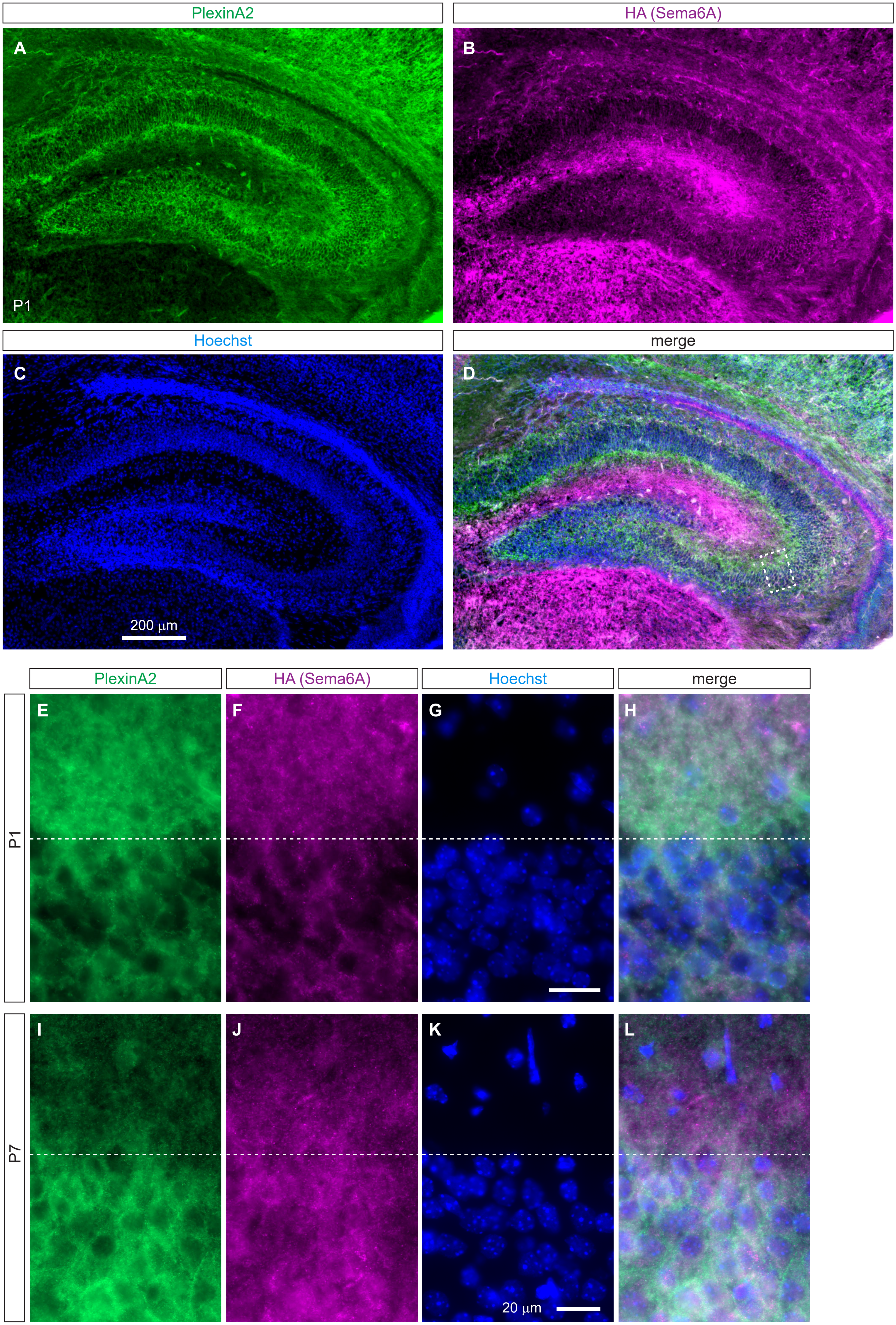
PlexinA2
HA (Sema6A)
A
B
P1
Hoechst
merge
C
D
200 µm
PlexinA2
HA (Sema6A)
Hoechst
merge
E
F
G
H
P1
I
J
K
L
P7
20 µm

**Figure S3.**

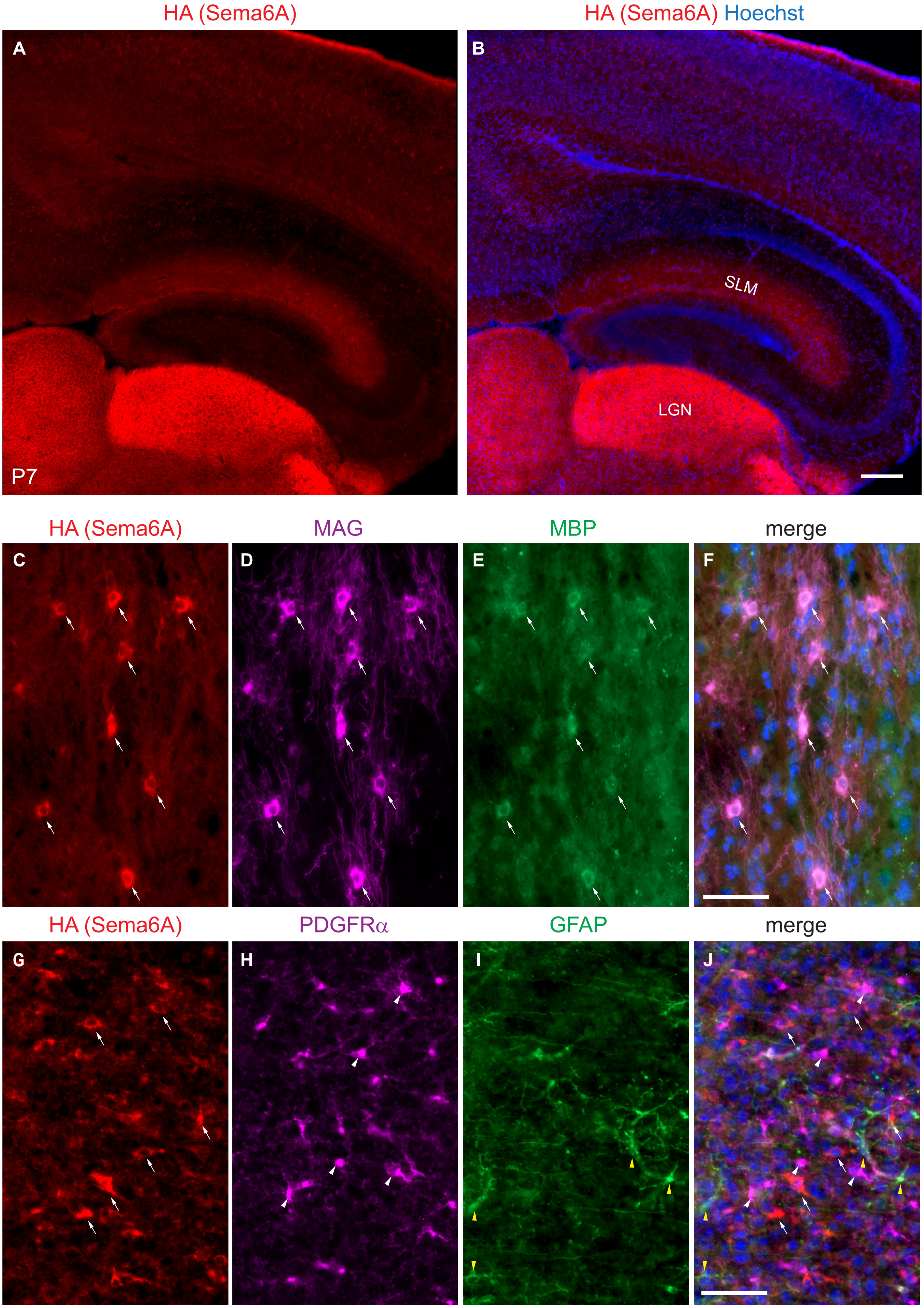
HA (Sema6A)
HA (Sema6A) Hoechst
A
B
P7
SLM
LGN
HA (Sema6A)
MAG
MBP
merge
C
D
E
F
HA (Sema6A)
PDGFRα
GFAP
merge
G
H
I
J

**Figure S4.**

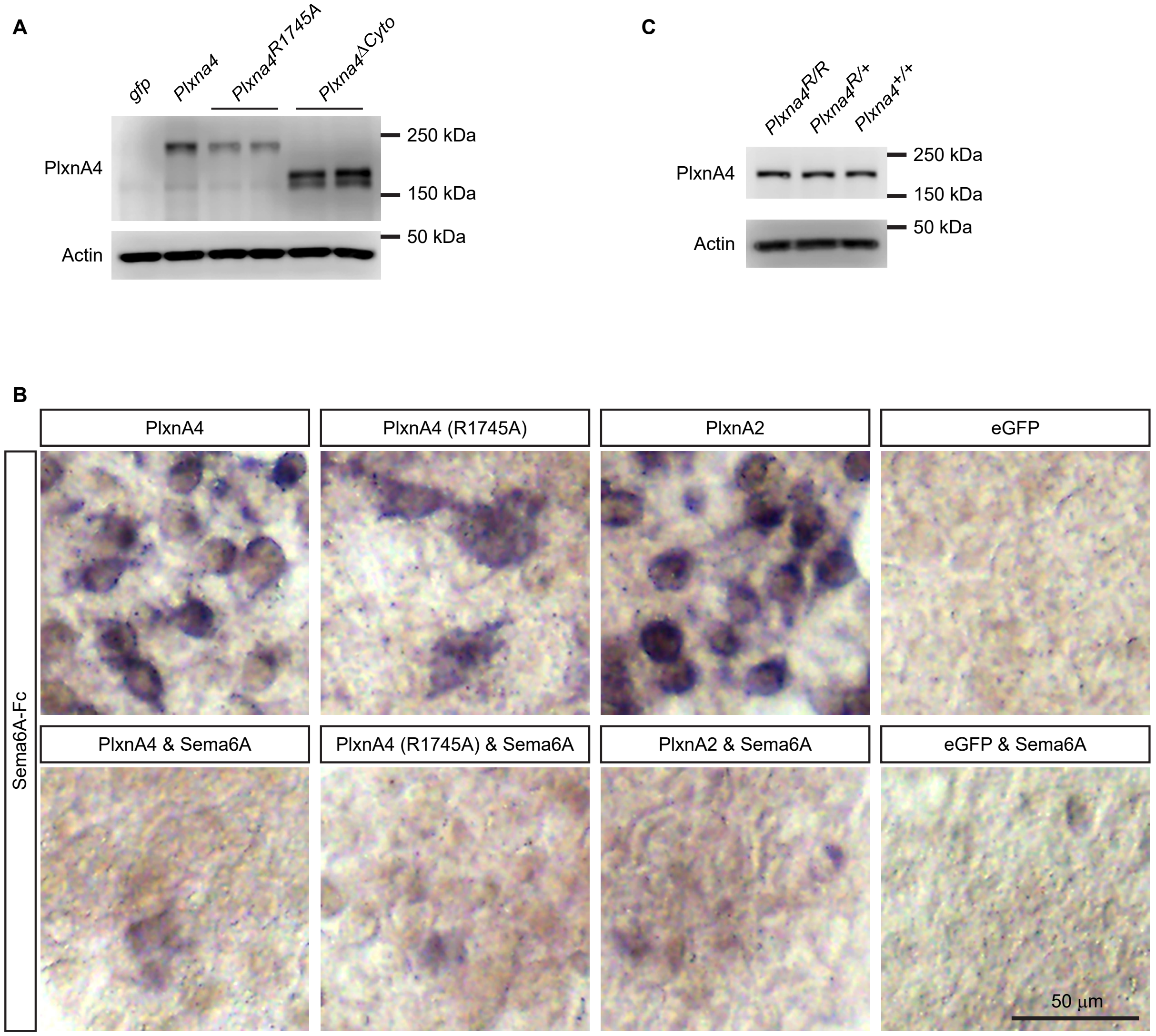
A
gfp
Plxna4
Plxna4R1745A
Plxna4ΔCyto
PlxnA4
Actin
250 kDa
150 kDa
50 kDa
C
Plxna4R/R
Plxna4R/+
Plxna4+/+
PlxnA4
Actin
250 kDa
150 kDa
50 kDa
B
PlxnA4
PlxnA4 (R1745A)
PlxnA2
eGFP
PlxnA4 & Sema6A
PlxnA4 (R1745A) & Sema6A
PlxnA2 & Sema6A
eGFP & Sema6A
Sema6A-Fc
50 µm

**Figure S5.**

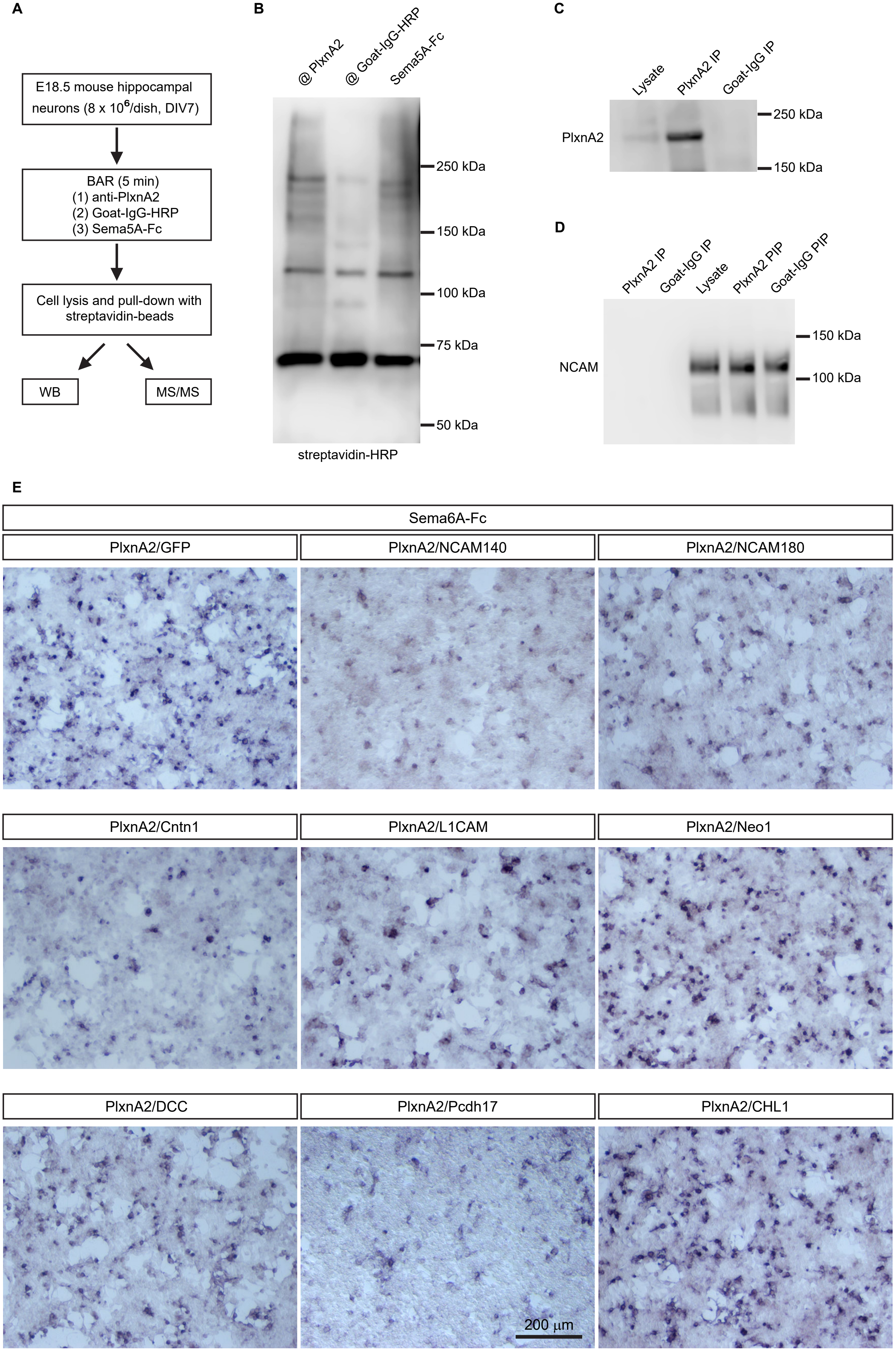
A
E18.5 mouse hippocampal neurons (8 x 10^6/dish, DIV7)
BAR (5 min)
(1) anti-PlxnA2
(2) Goat-IgG-HRP
(3) Sema5A-Fc
Cell lysis and pull-down with streptavidin-beads
WB
MS/MS
B
@ PlxnA2
@ Goat-IgG-HRP
Sema5A-Fc
250 kDa
150 kDa
100 kDa
75 kDa
50 kDa
streptavidin-HRP
C
Lysate
PlxnA2 IP
Goat-IgG IP
PlxnA2
250 kDa
150 kDa
D
PlxnA2 IP
Goat-IgG IP
Lysate
PlxnA2 PIP
Goat-IgG PIP
NCAM
150 kDa
100 kDa
E
Sema6A-Fc
PlxnA2/GFP
PlxnA2/NCAM140
PlxnA2/NCAM180
PlxnA2/Cntn1
PlxnA2/L1CAM
PlxnA2/Neo1
PlxnA2/DCC
PlxnA2/Pcdh17
PlxnA2/CHL1
200 μm

**Figure S6.**

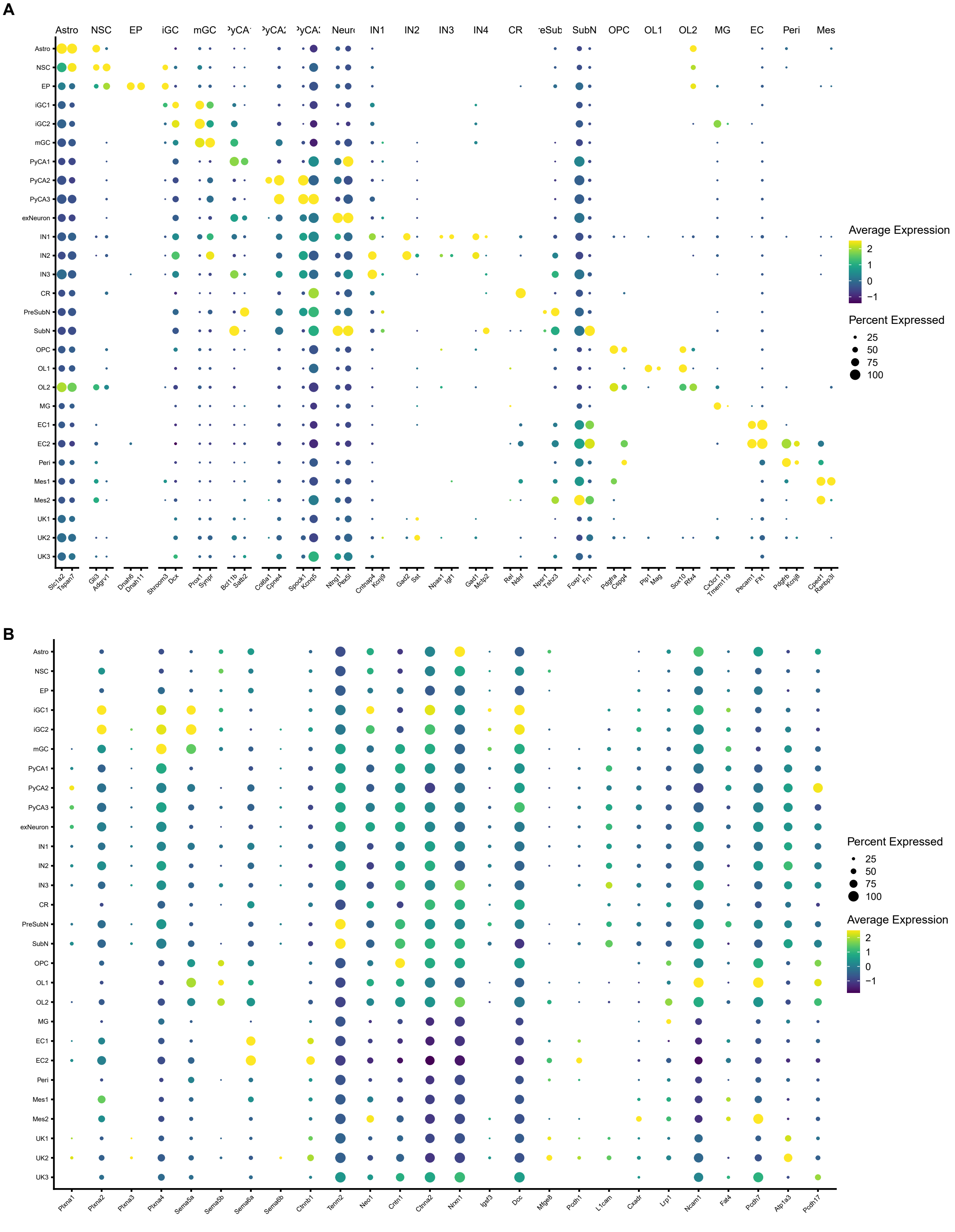

A
Astro
NSC
EP
iGC
mGC
IN1
IN2
IN3
IN4
CR
SubN
OPC
OL1
OL2
MG
EC
Peri
Astro
NSC
EP
iGC1
iGC2
mGC
PyCA1
PyCA2
PyCA3
exNeuron
IN1
IN2
IN3
CR
PreSubN
SubN
OPC
OL1
OL2
MG
EC1
EC2
Peri
Mes1
Mes2
UK1
UK2
UK3
Average Expression
Percent Expressed
B
Plxna1
Plxna2
Plxna3
Plxna4
Sema5a
Sema5b
Sema6a
Sema6b
Ctnnb1
Tenm2
Neo1
Cntn1
Ctnna2
Nrxn1
Igsf3
Dcc
Mfge8
Pcdh1
L1cam
Cxadr
Lrp1
Ncam1
Fat4
Pcdh7
Atp1a3
Pcdh17


**Figure S7.**

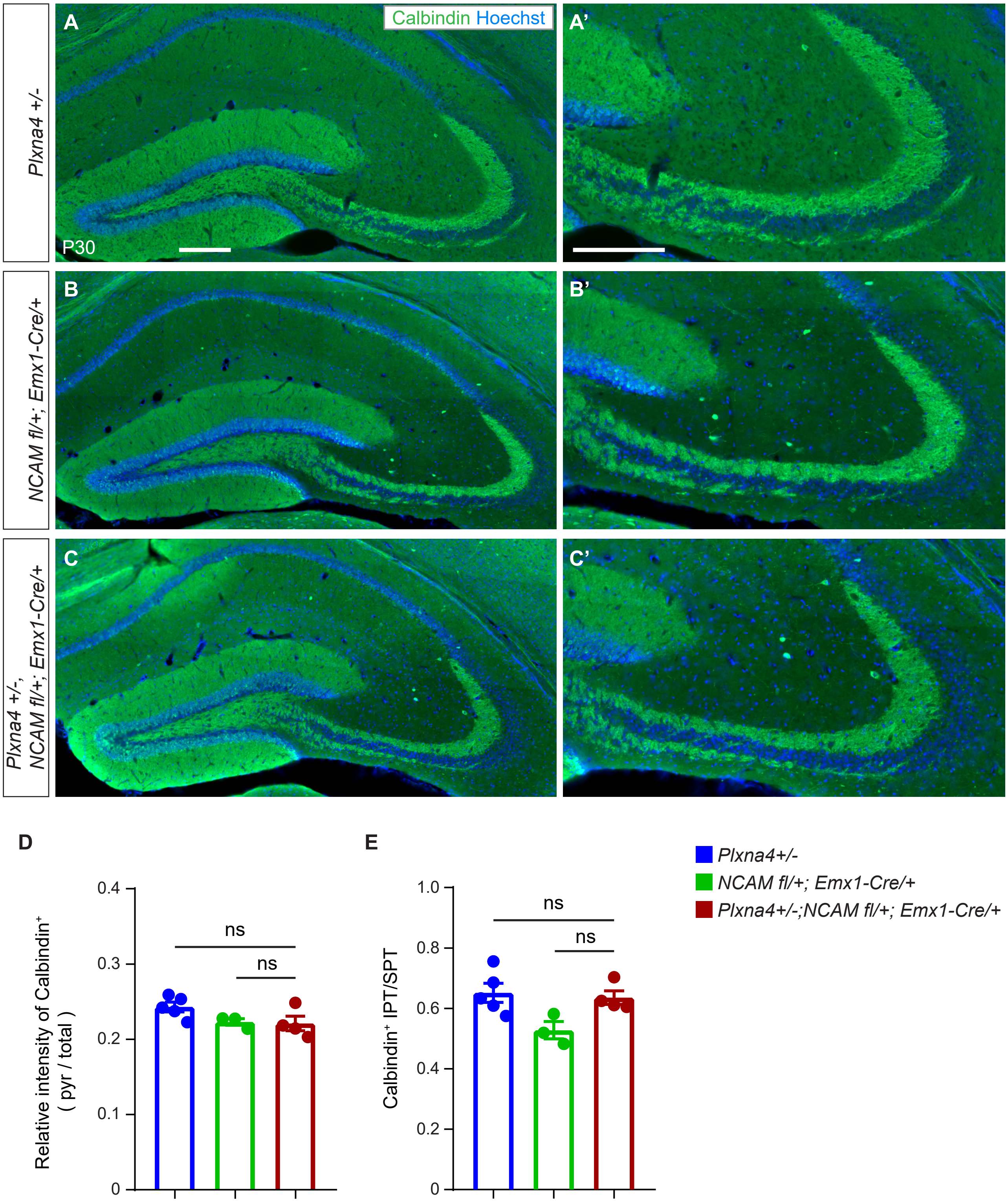

Calbindin Hoechst
A
A'
B
B'
C
C'
Plxna4 +/-
NCAM fl/+; Emx1-Cre/+
Plxna4 +/-,
NCAM fl/+; Emx1-Cre/+
P30
D
E
Relative intensity of Calbindin+
( pyr / total )
0.4
0.3
0.2
0.1
0
ns
ns
Calbindin+ IPT/SPT
1.0
0.8
0.6
0.4
0.2
0
ns
ns
Plxna4+/-
NCAM fl/+; Emx1-Cre/+
Plxna4+/-;NCAM fl/+; Emx1-Cre/+


**Figure S8.**

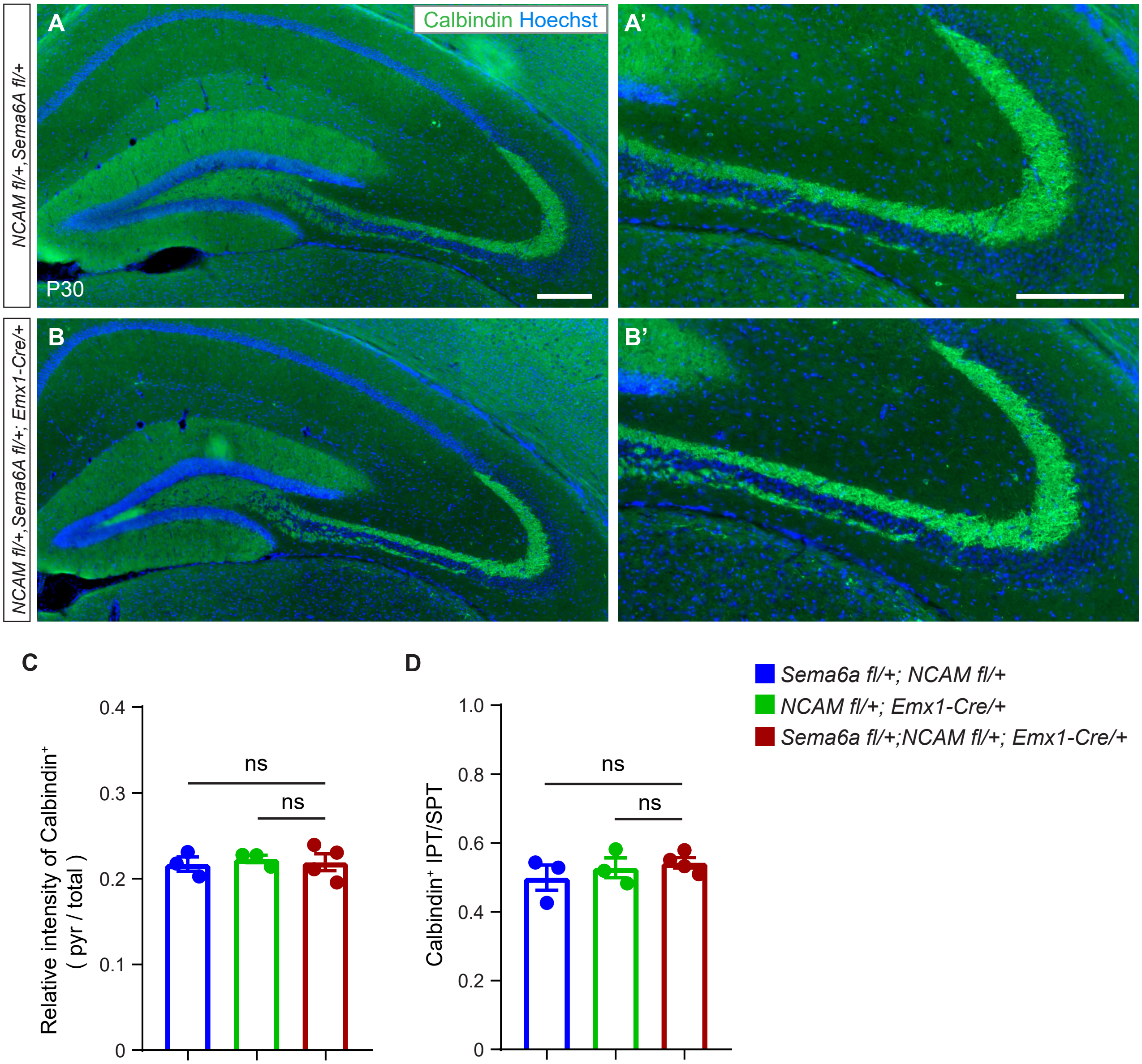

Calbindin Hoechst
A
A'
B
B'
P30
NCAM fl/+, Sema6A fl/+
NCAM fl/+, Sema6A fl/+; Emx1-Cre/+
C
D
Relative intensity of Calbindin+ ( pyr / total )
0.4
0.3
0.2
0.1
0
ns
ns
Calbindin+ IPT/SPT
1.0
0.8
0.6
0.4
0.2
0
ns
ns
Sema6a fl/+; NCAM fl/+
NCAM fl/+; Emx1-Cre/+
Sema6a fl/+;NCAM fl/+; Emx1-Cre/+


**Figure S9.**

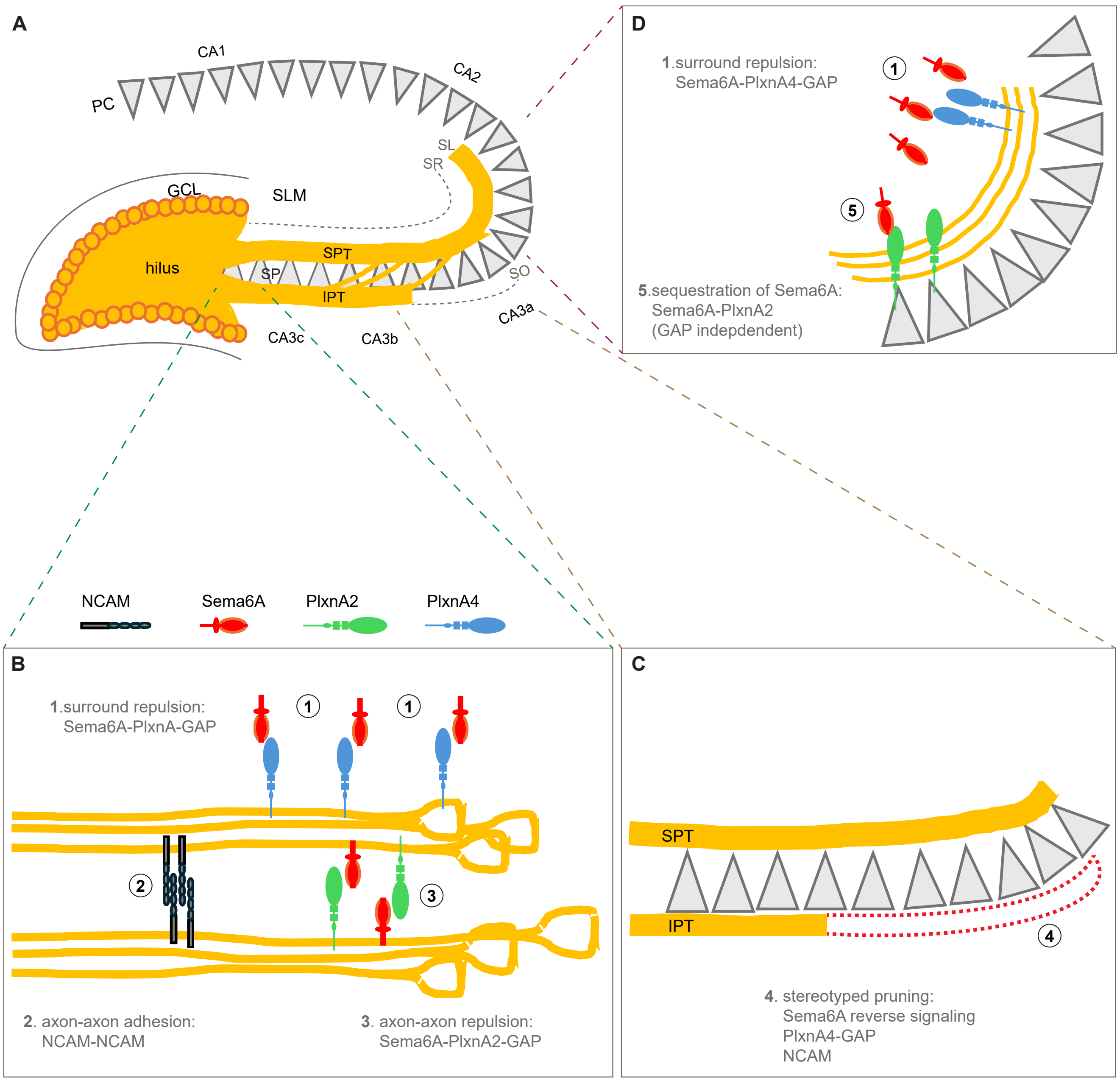

A
CA1
CA2
PC
SL
SR
GCL
SLM
hilus
SPT
SP
IPT
SO
CA3c
CA3b
CA3a
D
1.surround repulsion:
Sema6A-PlxnA4-GAP
1
5
5.sequestration of Sema6A:
Sema6A-PlxnA2
(GAP indepdendent)
NCAM
Sema6A
PlxnA2
PlxnA4
B
1.surround repulsion:
Sema6A-PlxnA-GAP
1
1
2
3
2. axon-axon adhesion:
NCAM-NCAM
3. axon-axon repulsion:
Sema6A-PlxnA2-GAP
C
SPT
IPT
4
4. stereotyped pruning:
Sema6A reverse signaling
PlxnA4-GAP
NCAM


**Figure S10.**
